# Subtelomeric regions and a repeat-rich chromosome harbor multicopy effector gene clusters with variable conservation in multiple plant pathogenic *Colletotrichum* species

**DOI:** 10.1101/2020.04.28.061093

**Authors:** Pamela Gan, Ryoko Hiroyama, Ayako Tsushima, Sachiko Masuda, Arisa Shibata, Akiko Ueno, Naoyoshi Kumakura, Mari Narusaka, Trinh Xuan Hoat, Yoshihiro Narusaka, Yoshitaka Takano, Ken Shirasu

**Affiliations:** RIKEN Center for Sustainable Resource Sciences, Yokohama, Kanagawa, Japan; John Innes Center, Norwich Research Park, Norwich, United Kingdom; Research Institute for Biological Sciences, Okayama Prefectural Technology Center for Agriculture, Forestry, and Fisheries, Okayama, Japan; Plant Protection Research Institute, Ha Noi city, Viet Nam; Graduate School of Agriculture, Kyoto University, Kyoto, Japan; Graduate School of Science, The University of Tokyo, Bunkyo, Tokyo, Japan

## Abstract

Members of the *Colletotrichum gloeosporioides* species complex are causal agents of anthracnose in a wide range of commercially important plants. To provide an in-depth overview of its diversity, we sequenced the genomes of fungi belonging to this group, including multiple strains of *C. fructicola* (*Cf*) and *C. siamense* (*Cs*), as well as representatives of three previously unsequenced species, *C. aenigma* (*Ca*), *C. tropicale* and *C. viniferum*. Comparisons between multiple *Cf* and *Cs* strains led to the identification of accessory regions that show variable conservation in both lineages. These accessory regions encode effector candidate genes, including homologs of previously characterized effectors, organized in clusters of conserved synteny with copy number variations in different strains of *Cf, Cs* and *Ca*. Analysis of highly contiguous assemblies of *Cf, Cs* and *Ca* strains revealed the association of such accessory effector gene clusters with subtelomeric regions and repeat-rich minichromosomes and provided evidence of gene transfer between these two genomic compartments. In addition, expression analysis indicated that paralogs associated with clusters of conserved synteny showed a tendency for correlated gene expression. These data highlight the importance of subtelomeric regions and repeat-rich chromosomes to the genome plasticity of *Colletotrichum* fungi.

## Introduction

Fungi within the genus *Colletotrichum* can be subdivided into species complexes consisting of closely related species (Cannon et al. 2012). Among them, members of the *Colletotrichum gloeosporioides* species complex (CGSC) are pathogens that cause significant damage to a wide range of commercially important plants (Weir et al. 2012). For example, *Colletotrichum fructicola* (*Cf*) infects avocado, apple, pear, strawberry, lemons, cocoa, coffee and yam (Weir et al. 2012). Further, different CGSC species have geographic and host range overlaps. For instance, *Cf, Colletotrichum siamense* (*Cs*), and *Colletotrichum aenigma* (*Ca*) have been identified as causal agents of strawberry anthracnose in Japan (Gan, Nakata, et al. 2016). Genus-wide genome comparisons between members of different species complexes have indicated that CGSC members have expanded their pathogenicity-associated genes including proteases, carbohydrate active enzymes and secondary metabolite (SM) biosynthesis genes compared to other *Colletotrichum* (Baroncelli et al. 2016; Gan, Narusaka, et al. 2016), which may contribute to their role as broad host range pathogens.

Another class of pathogenicity-associated genes are those encoding effectors, small, secreted proteins that are hypothesized to contribute to infection by manipulation of host cell structure and function (Kamoun 2006). In turn, host plants have evolved to recognize these proteins, resulting in host cell death and resistance (Jones and Dangl 2006; Dodds and Rathjen 2010). Given this scenario, effectors are often under a diversifying selection to evade recognition by hosts. An emerging paradigm is that the need to balance diversifying selection with maintenance of housekeeping genes has driven the compartmentalization of pathogen genomes into fast-evolving genomic regions, encoding effector genes, and conserved regions, encoding housekeeping genes (Dong et al. 2015).

Chromosome-level assemblies of *Colletotrichum lentis* and *Colletotrichum higginsianum*, which belong to the *Colletotrichum destructivum* species complex, have revealed fast-evolving genomic regions in repeat-rich, pathogenicity-associated minichromosomes (Dallery et al. 2017; Plaumann et al. 2018; Bhadauria et al. 2019). Further, the genome of *Colletotrichum orbiculare* 104-T from the *C. orbiculare* species complex, which lacks minichromosomes (Taga et al. 2015), shows signatures of compartmentalization with distinct, gene-poor, AT-rich, transposable element (TE)-dense regions, as well as gene-rich, GC-rich, TE-poor, regions (Gan et al. 2013). Similar TE-dense, AT-rich compartments in other plant pathogenic fungi are hypothesized to have undergone repeat-induced point (RIP) mutations, a fungal defense mechanism against TEs that occurs during sexual reproduction, leading to effector gene diversification (Rouxel et al. 2011).

This study aims to gain insights into the evolution of CGSC genomes by comparing multiple strains from different species with shared geographical and/or host ranges. Specifically, we aimed to identify fast-evolving, non-conserved genomic regions within the species complex. By analyzing the location of identified accessory genomic regions with variable conservation patterns, we gained insights into the mechanisms by which pathogen genomes evolve, acquire and organize pathogenicity-related genes.

## Results

### High quality genome assemblies of *C. fructicola, C. siamense* and *C. aenigma*

The genomes of 14 CGSC strains showing differences in pathogenicity in the field were selected for sequencing. The pathogenicity of these strains were tested against *Fragaria vesca* and *Fragaria* × *ananassa* var. Sachinoka (Supplementary Figs. S1-2). Based on these results, pathogenicity on strawberries is not determined by species, with *Cf* strains Nara gc5, S1 and S4 causing severe disease symptoms on both hosts, while strains Cf413, Cf245 and Cf415 did not. Similarly, *Cs* Cg363, CAD1 and CAD5 caused lesions on both hosts while CAD2 and CAD4 did not. *Cf* Nara gc5, *Cs* Cg363 and *Ca* Cg56 were selected for PacBio sequencing as they cause disease symptoms on strawberry leaves despite belonging to different species according to genome-wide SNP analysis (Fig. 1, Supplementary Fig. S3). In addition, *Cf* Cf413 was also sequenced by PacBio as a representative non-pathogenic strain.

**Figure 1.**
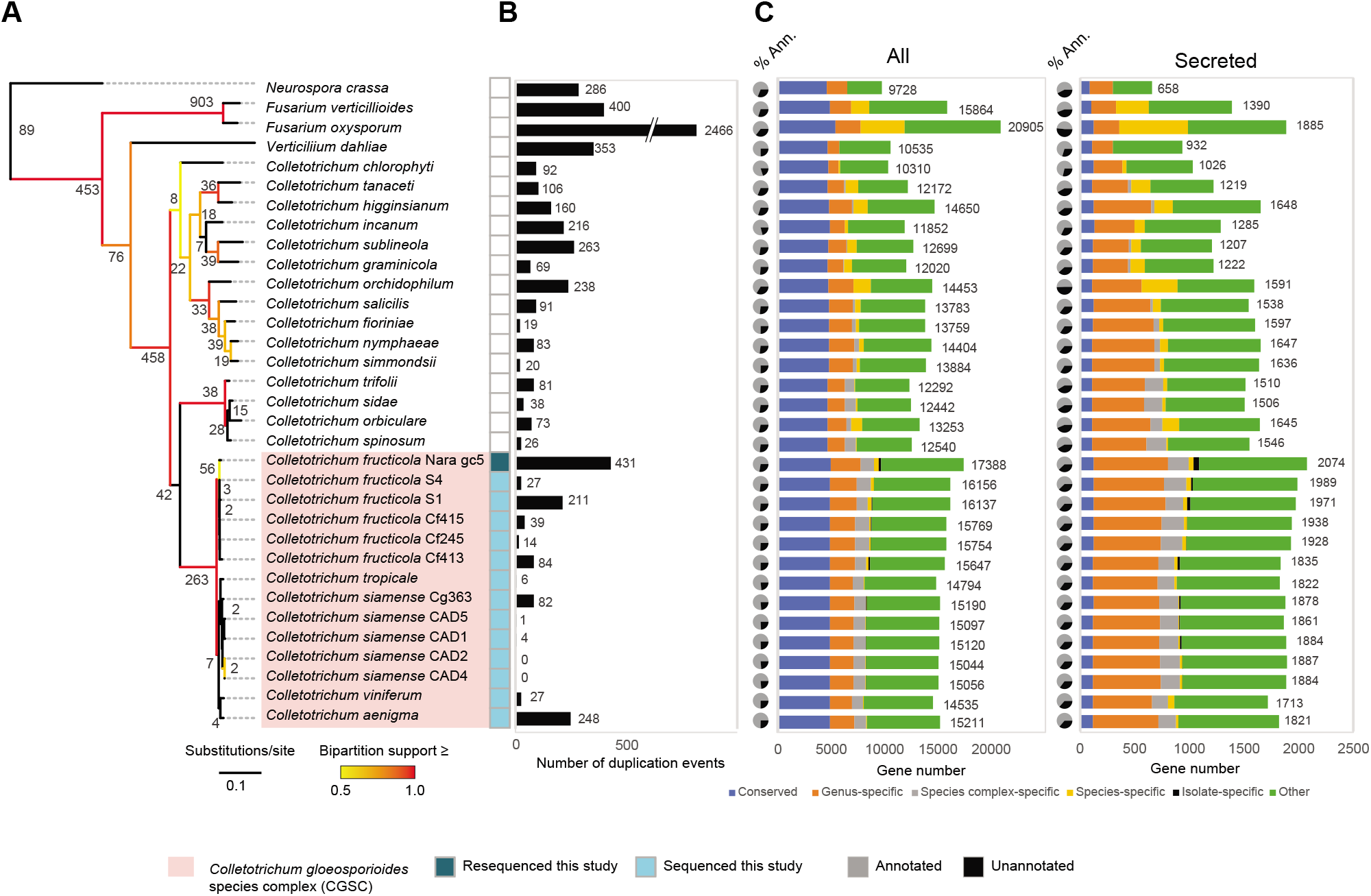
Relationship between sequences in the 14 sequenced *Colletotrichum gloeosporioides* species complex isolates. (A) Phylogenetic relationship between 33 different ascomycetes including 29 members from the *Colletotrichum* genus based on all orthogroups identified in OrthoFinder. Numbers at the nodes represent the number of estimated duplication events occurring at each node with at least 50 % support according to OrthoFinder analysis. (B) Bars representing numbers of OrthoFinder-estimated strain-specific duplication events with at least 50 % support. (C) Conservation pattern of all genes in each sequenced strain. (D) Conservation pattern of secreted genes by strain. Pie charts indicate percentage of genes that could be annotated with an InterPro domain. Gene numbers are shown beside bars.

All 14 genome assemblies were estimated to include at least 99.3 % of the gene coding space (Table 1). Genomes of the 14 CGSC strains were annotated and the conservation of genes was determined using OrthoFinder (Fig. 1). The number of predicted genes in these CGSC strains ranges from 14,535 to 17,388, which is generally higher than other *Colletotrichum* species (Fig. 1C). Greater numbers of carbohydrate active enzymes (CAzymes), secreted proteases and secondary metabolite (SM)-encoding genes were detected in all CGSC strains relative to non-CGSC *Colletotrichum* spp. regardless of pathogenicity, indicating that the expansion of these classes of genes is a general feature of this group (Supplementary Fig. S4). Further, only a limited number of strain-specific genes and duplication events were identified in the CGSC strains (Fig. 1A-C).

**Table 1.** Statistics of genomes assembled in this study. The completeness of the genome assemblies was assessed by estimating the conservation of BUSCO *Pezizomycotina* conserved genes.

| Species | Strain | Assembly size | Scaffold number | N50 (bp) | L50 | % BUSCO genes |  | Predicted genes |
| --- | --- | --- | --- | --- | --- | --- | --- | --- |
|  |  |  |  |  |  | Complete | Fragmented |  |
| *C. fructicola* | Nara gc5 | 59.6 Mb | 13 | 5,461,344 | 5 | 99.5 | 0.4 | 17,388 |
|  | Cf413 | 56.5 Mb | 14 | 4,902,990 | 5 | 99.6 | 0.4 | 15,647 |
|  | S1 | 59.0 Mb | 600 | 1,273,051 | 16 | 99.4 | 0.3 | 16,137 |
|  | CfS4 | 57.2 Mb | 1,588 | 306,788 | 52 | 99.4 | 0.5 | 16,156 |
|  | Cf245 | 56.1 Mb | 1,194 | 400,477 | 46 | 99.5 | 0.4 | 15,754 |
|  | Cf415 | 56.0 Mb | 805 | 413,455 | 41 | 99.4 | 0.5 | 15,769 |
| *C. siamense* | Cg363 | 62.9 Mb | 22 | 5,441,060 | 5 | 99.5 | 0.3 | 15,190 |
|  | CAD1 | 58.4 Mb | 311 | 881,809 | 22 | 99.6 | 0.3 | 15,120 |
|  | CAD2 | 58.1 Mb | 214 | 709,060 | 25 | 99.5 | 0.5 | 15,044 |
|  | CAD4 | 58.2 Mb | 199 | 823,408 | 24 | 99.6 | 0.3 | 15,056 |
|  | CAD5 | 57.6 Mb | 294 | 656,815 | 27 | 99.6 | 0.3 | 15,097 |
| *C. aenigma* | Cg56 | 59.2 Mb | 79 | 5,181,253 | 5 | 99.7 | 0.3 | 15,211 |
| *C. viniferum* | CgW01 | 68.5 Mb | 5,864 | 146,824 | 138 | 99.3 | 0.6 | 14,535 |
| *C. tropicale* | S9275 | 55.8 Mb | 789 | 691,924 | 24 | 99.4 | 0.4 | 14,794 |

Identification of the telomeric repeat TTAGGG revealed that the PacBio-sequenced genomes of *Cf* Nara gc5, Cf413 and *Cs* Cg363 each possess 10, 10, and 7 contigs of greater than 100 kb enriched with telomeric repeats at both ends (≥25 copies TTAGGG/terminal 10 kb), suggesting that each of these assemblies include 10, 10 and 7 complete telomere-to-telomere chromosomes respectively (Fig. 2A). The *Cf* Nara gc5 assembly is the most contiguous, consisting of 13 contigs: 12 of which are greater than 100 kb, and the remaining contig being homologous to the previously sequenced *Cf* mitochondrial genome (KX034082; Liang et al. 2017) (Supplementary Fig. S3B).

**Figure 2.**
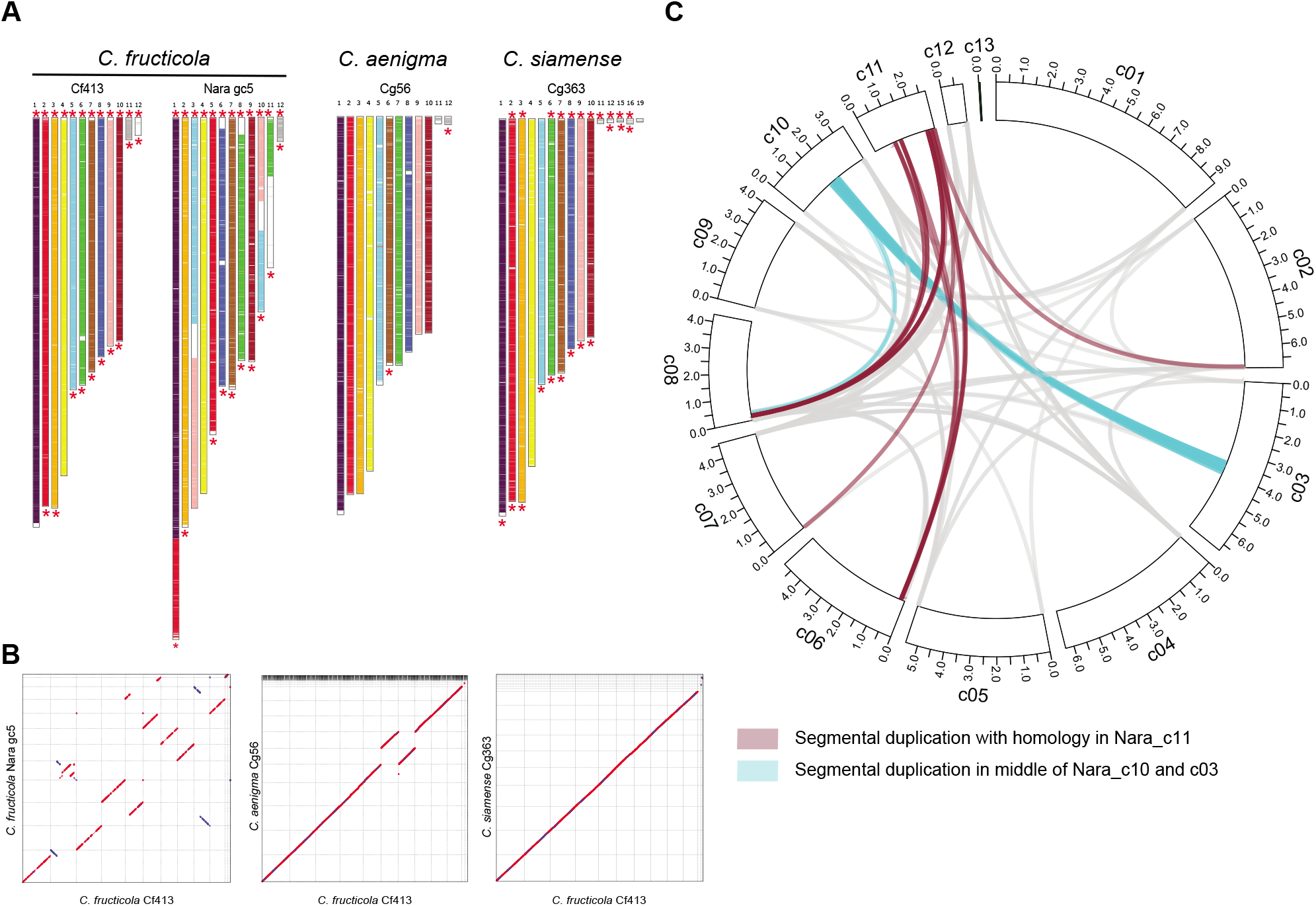
Genome rearrangements in the *Colletotrichum gloeosporioides* species complex. (A) Contigs are colored according to the contig of homology in Cf413. White regions are regions without synteny in Cf413. (B) Dot plot representing forward (red) and reverse (blue) hits with conservation between Nara gc5, Cg363 and Cg56 against Cf413. Hits were identified using nucmer with the maxmatch settings. Only hits of greater than 10 kb are shown. (C) Within the *C. fructicola* Nara gc5 genome, segmental duplications were detected mostly between the ends of contigs and with the repeat-rich chromosome Nara_c11. An additional segmental duplication was detected between the middle of Nara_c03 and c10 in a region corresponding to a potential chromosomal breakpoint. Ticks represent 0.5 Mb.

Whole genome alignments between *Cf* Nara gc5, Cf413, *Ca* Cg56 and *Cs* Cg363, were performed to detect the presence of genomic rearrangements. Despite their close phylogenetic relationship (Supplementary Fig. S3A), multiple rearrangements were detected in *Cf* Nara gc5 relative to Cf413 (Fig. 2A-B). In contrast, Cf413 shares a high degree of collinearity with the more distantly related strains, *Cs* Cg363 and *Ca* Cg56. Using *Cs* Cg363 as a reference outgroup, Nara_c01, c05, c03, c10, c08 and c11 appear to have resulted in novel chromosomes derived from translocations (Fig. 2A-B). To confirm whether these rearrangements are *Cf* Nara gc5-specific, mapping coverages of PacBio reads and 500 bp insert-sized, paired-end reads were visually inspected around 14 rearrangement sites representing synteny breaks of greater than 100 kb between Nara gc5 and Cf413 identified by nucmer (Supplementary Fig. S5). From this analysis, 3 and 5 of 14 of these rearrangements were also identified in *Cf* strains S1 and S4, respectively (Supplementary Fig. S5). Read mapping analysis also revealed that approximately 1.2 Mb encompassing 400 genes in Cf413 including 20 candidate effectors and two secondary metabolite clusters on one arm of 413_c09 is duplicated, as the read mapping rate is approximately 2-fold in this region compared to the rest of the genome (Supplementary Fig. S6). This duplication appears to be strain-specific since the only *Cf* strain showing this pattern was Cf413.

Segmental duplications in *Cf* Nara gc5 and Cf413 were also identified by whole genome alignments within each genome. In both strains, the majority of intercontig segmental duplications of greater than 10 kb were detected between the ends of contigs (Fig. 2C, Supplementary Fig. S6A). Segmental duplications were also identified at points of large scale structural variations, including between the region just upstream of the 413_c09 duplication and the end of 413_c01 (Supplementary Fig. S6), as well as between Nara_c03 and Nara_c10, where the existence of a chromosomal translocation is predicted (Fig. 2C, Supplementary Fig. S7).

### CGSC members differ in LTR transposable element abundances

Transposable elements (TEs) are known to influence genomic landscapes leading to the compartmentalization and diversification of genomes (Dong et al. 2015). Thus, we investigated the composition of repeat elements in the genomes studied. Analysis of the repeat contents of the PacBio-sequenced CGSC strains and the non-CGSC clade member *C. higginsianum* IMI 349063 revealed that retrotransposons, especially LTR retrotransposons, are more abundant in CGSC strains compared to DNA transposons, in contrast to *C. higginsianum* (Fig. 3A). However, retrotransposon content varied between strains with LINE-type repeats showing the greatest variation, ranging from 0.08 % of the *Cs* Cg363 genome to 0.44 % of the *Cf* Nara gc5 and *Ca* Cg56 genomes, a difference greater than 5-fold. Repeat element composition showed considerable variation even between different strains of the same species with *Cf* Nara gc5 having 2.3 times more *Gypsy*-type LTR retrotransposons as a proportion of the total genome size compared to Cf413. Genome sizes and gene numbers were not found to be related to repeat abundance (Supplementary Fig. S8A-B).

**Figure 3.**
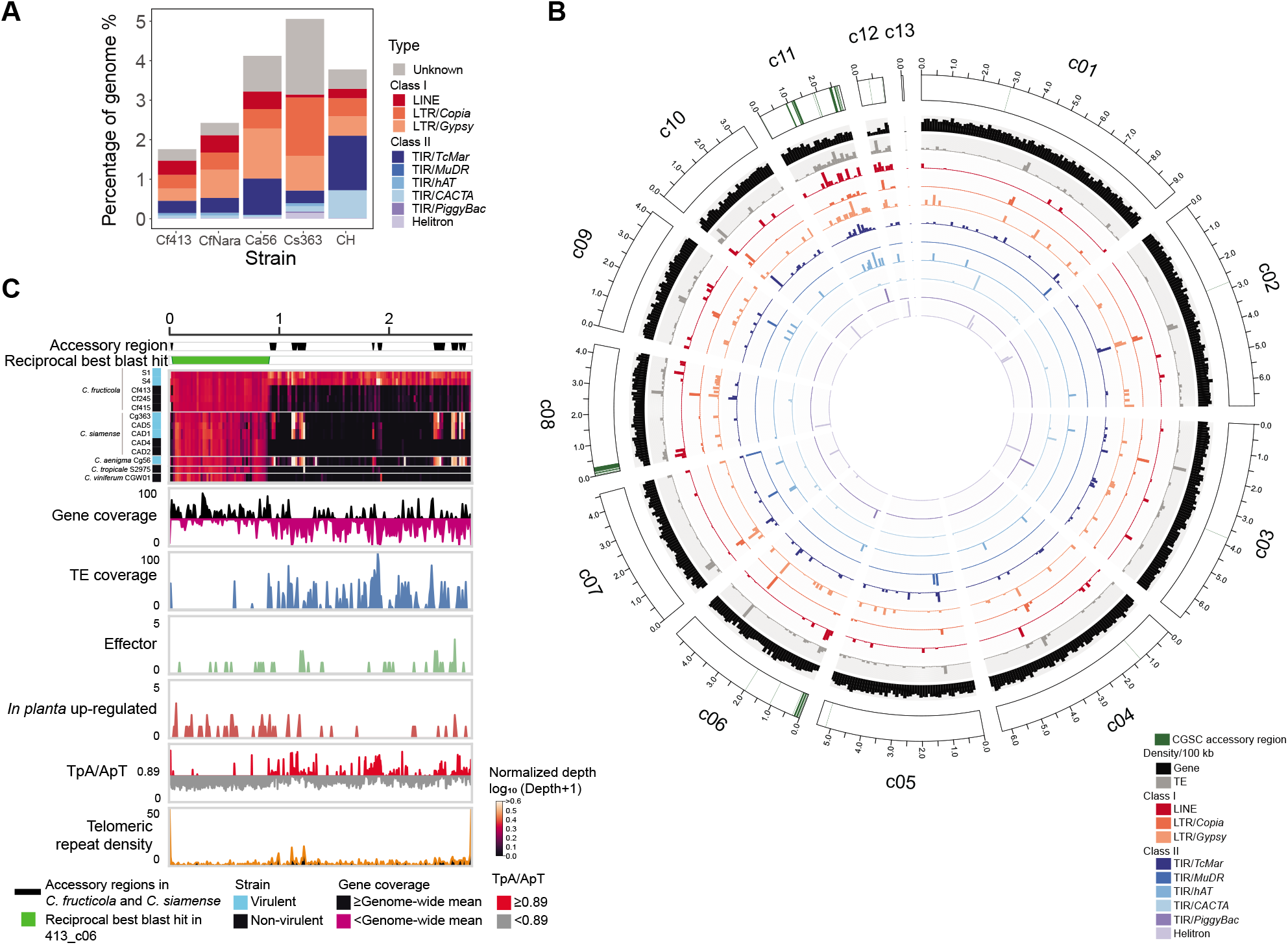
Repeats in *Colletotrichum gloeosporioides* species complex members. (A) Proportions of sequences in the genomes of *C. fructicola* Nara gc5 (CfNara) and Cf413, *C. aenigma* Cg56 (Ca56), *C. siamense* Cg363 (Cs363) and *C. higginsianum* IMI 349063 (CH) associated with TEs of the major superfamilies. (B) Distribution of TEs in *C. fructicola* Nara gc5. Ticks represent 0.5 Mb. (C) Compartmentalization of Nara_c11 into TE-rich, gene-poor regions with CGSC accessory regions and TE-poor, gene-dense regions. Nara_c11 shows enrichment of telomeric repeats at both ends suggesting it is a complete chromosome. The number of effectors, in planta up-regulated genes and telomeric repeats were calculated in 10 kb windows.

To identify potential repeat compartmentalization, the distributions of repeats were characterized in greater detail. In the two *Cf* strains, LINE-type TEs were enriched at the ends of contigs and in smaller contigs (Fig. 3B, Supplementary Fig. S8C). Of note, Nara_c11 has distinct compartmentalization with one half that is TE-rich and gene-sparse, while the other half is TE-sparse and gene-rich (Fig. 3C). The TE-rich region of Nara_c11 also has higher TpA/ApT ratios relative to the rest of the chromosome (Fig. 3C), which is a signature of repeat-induced point mutation.

### Accessory regions with variable conservation are associated with TEs and subtelomeric regions in different CGSC lineages

To identify rapidly evolving genomic compartments, reads from the sequenced CGSC strains were mapped to the *Cf* Nara gc5 assembly. Reads from all sequenced strains mapped to 83.75 % of the Nara gc5 assembly (49.91 Mb; log_10_ (normalized depth+1) ≥ 0.1) indicating that most of the genome is conserved (Supplementary Fig. S9). In addition, 0.82 Mb (1.38 %) of the genome was conserved in all *Cf* strains, but not in strains from other species, potentially representing *Cf-*specific regions. Also, 3.67 Mb of the assembly was detected as dispensable, being present in at least one *Cf* strain, but not in all of them. These dispensable regions overlapped with repeat-rich regions at the ends of Nara_c08 and Nara_c06, as well as the repeat-rich region of Nara_c11 (Supplementary Fig. S9, Fig. 3B).

The dispensable region of Nara_c11 (from position 910,000 to the end of Nara_c11) is TE-rich and gene-poor (mean TE coverage of 14.32%, median gene coverage of 34.11 %, median TpA/ApT ratio of 0.77, mean number of *in planta* up-regulated genes of 0.12 genes/10 kb) (Supplementary Table S1). This was distinct from the rest of the contig (mean TE coverage of 1.88%, median gene coverage of 52.95 %, median TpA/ApT ratio of 0.60, mean number of *in planta* up-regulated genes of 0.40 genes/10 kb), that shared homology to 413_c06. Instead, the dispensable region of Nara_c11 was more similar to Nara_c12, which encodes a potential minichromosome conserved in all other *Cf* strains according to read mapping analysis (mean TE coverage of 12.1%, median gene coverage of 17.0 %, median TpA/ApT ratio of 0.65, Supplementary Table S1). However, the mean number of *in planta* up-regulated genes in Nara_c12 is lower than that of the Nara_c11 dispensable region (0.01 genes/10 kb).

Interestingly, read mapping analysis revealed that 0.84 Mb of *Cf* Nara gc5 dispensable regions also have variable conservation in *Cs* strains (present in at least one *Cs* strain but not all), indicating the existence of sequences that are conserved in only subsets of *Cf* and *Cs* strains. We refer to these *Cf* and *Cs* variably conserved regions as CGSC accessory regions since they are dispensable in multiple CGSC lineages. In *Cf* Nara gc5, CGSC accessory regions significantly associate with sequences enriched with the telomeric repeat TTAGGG (Fig. 4A), indicating that they are present in subtelomeres. In *Cf* Nara gc5, LINE and *Copia* TEs also significantly overlapped with CGSC accessory regions (P<0.05, Fig. 4A, Supplementary Table S2). In contrast, only a 0.14% (0.08 Mb) of Cf413 fits the same criteria (present according to read mapping depth in some but not all *Cf* and *Cs* strains), but no TEs were found to overlap these regions. However, in the pathogenic *Cs* strain, Cg363, 0.66 Mb CGSC accessory regions (present in some but not all *Cf* and *Cs* strains) were identified. In Cg363, these regions significantly overlap with subtelomeric regions, *Copia, Gypsy* and *TcMar-type* TEs (P<0.05, Supplementary Table S2).

**Figure 4.**
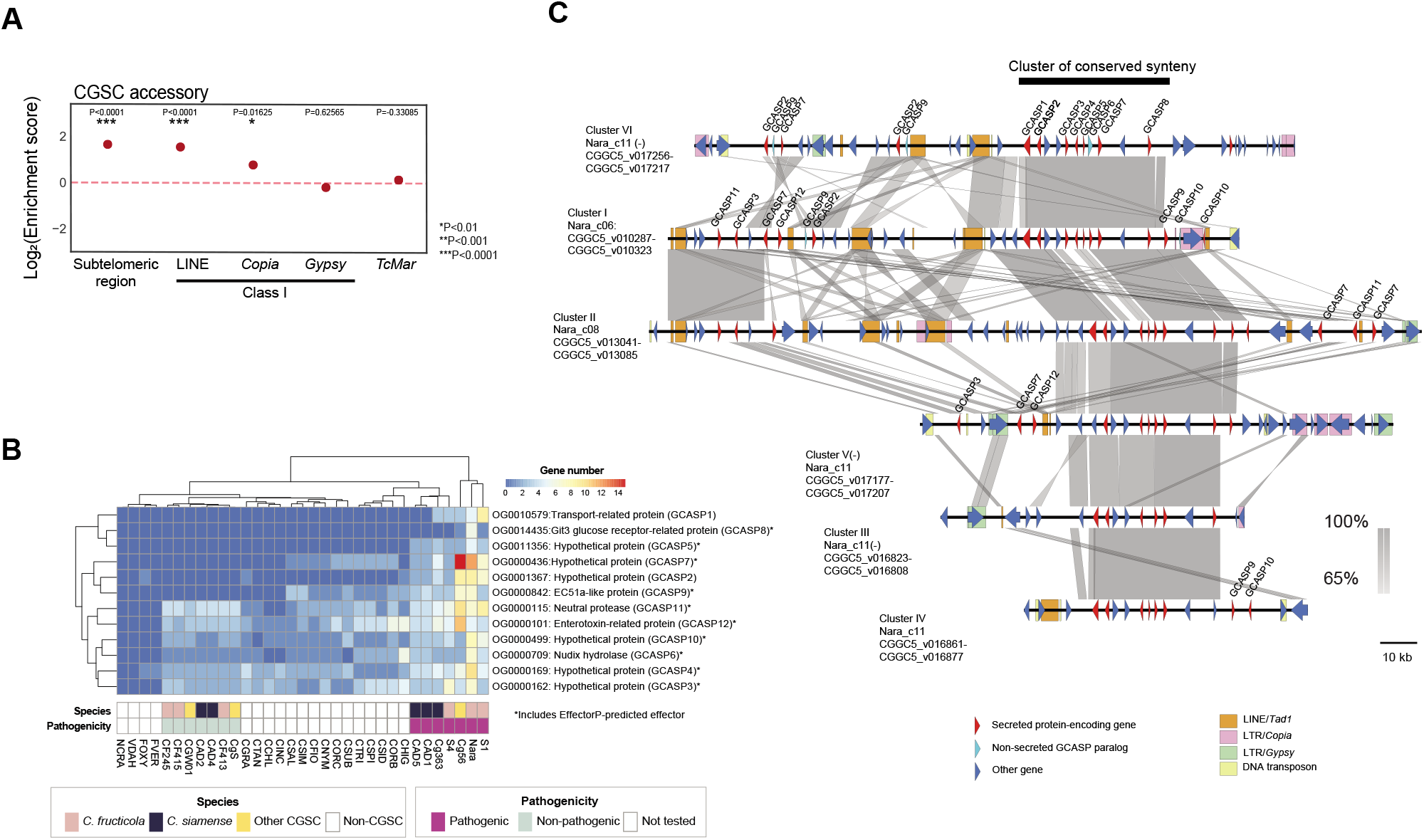
CGSC accessory regions. (A) Association of specific features with CGSC accessory regions. Enrichment is the number of observed overlaps between CGSC accessory regions with specific features of interest normalized by the expected number of overlaps if features of interest were distributed at random within each chromosome (median of 10,000 simulated replicates). A log_2_(Enrichment score) of 0 indicates no difference compared to the randomized simulations. (B) Conservation of orthogroups in 33 analyzed Ascomycetes with at least one member in the CGSC accessory regions and one predicted extracellular protein. Strain abbreviations are fully defined in Supplementary Table S5. (C) Segmental duplications of accessory regions encoding GCASPs in Colletotrichum fructicola Nara gc5. Ribbons indicate regions of homology of greater than 1 kb and BLASTn E-values <0.0001.

A total of 205 genes were identified in *Cf* Nara gc5 CGSC accessory regions. Proteins encoded by these genes were assigned to 80 orthogroups by OrthoFinder analysis, except for 5 proteins, with no known homologs amongst the 32 other analyzed fungal strains. Of these 80 orthogroups, 16 include at least one predicted Nara gc5 extracellular protein. Further, 12 of these 16 orthogroups encode more than one Nara gc5 paralog (Fig. 4B). Interestingly, based on the copy number profiles of these 12 orthogroups, the 7 non-pathogenic CGSC strains clustered apart from 7 pathogenic strains, irrespective of their species of origin (Fig. 4B). For ease of reference, these 12 orthogroups will be referred to as **<u>G</u>**loeosporioides species **<u>C</u>**omplex **<u>A</u>**ccessory **<u>S</u>**ecreted **<u>P</u>**aralog (GCASP) groups in the rest of the text.

Of the 12 GCASP orthogroups, 7 have no known function, including one group of *C. higginsianum* effector candidate EC51a homologs (Fig. 4B). On the other hand, GCASPs with homologs of known function include orthologs of CtNudix, a previously characterized nudix hydrolase effector from *C. lentis* (Bhadauria et al. 2012), Git3 glucose-receptor-related proteins, enterotoxin-related proteins, and proteases (Fig. 4B). Furthermore, 10 of the 12 GCASP orthogroups include at least one EffectorP-predicted effector.

### CGSC accessory regions harbor effector candidate clusters that are segmentally duplicated in *Cf* Nara gc5

In *Cf* Nara gc5, GCASP genes are organized in 6 paralogous gene clusters of conserved synteny, with one cluster each in subtelomeric regions of Nara_c06 and c08 and two pairs of tandemly duplicated clusters in Nara_c11 (Cluster I-VI in Fig. 4c, Supplementary Fig. S10, Supplementary Table S3). TEs associated with GCASP clusters of conserved synteny were also examined in *Cf* Nara gc5. All 6 clusters were located close to LINE/*Tad1*-type retrotransposons (Fig. 4C), including sequences with homology to CgT1, a *C. gloeosporioides* biotype-specific TE (He et al. 1996). In addition, *Copia* and *Gypsy*-type LTR retrotransposons including reverse transcriptase-coding sequences were identified flanking Clusters I, III, and V (Fig. 4C).

For insight into GCASP evolution, the phylogenies and locations of these genes in *Cf* Nara gc5 were examined. GCASPs 1-10 have paralogs in at least 2 of the 6 accessory clusters of conserved synteny, with GCASPs 1, 5 and 8 found only in these clusters. On the other hand, GCASPs 2, 7 and 9 also have paralogs in CGSC accessory regions outside of these clusters (Fig. 4C). Further, GCASPs 3, 4, 6 and 10 have paralogs in CGSC accessory regions as well as in the core genome (Fig. 4C, Supplementary Fig. S11). In contrast to GCASPs 1-10, members of GCASPs 11 and 12 are in core and CGSC accessory regions, but not in the 6 accessory clusters. For the 5 GCASPs with homologs in core regions (except GCASP3), *Cf* Nara gc5 accessory paralogs form monophyletic clades with sequences from *Cf* S1, S4, *Cs* Cg363, CAD1, CAD5 and *Ca* Cg56, which are separate from clades of core paralogs, indicating a common origin of these sequences in the 3 species (Supplementary Fig. S11). However, *Cs* accessory GCASPs tend to show greater sequence divergence from *Cf* compared to *Ca* sequences, consistent with the relatedness of the three species. Accessory GCASP10 sequences are absent from *Ca* Cg56, while accessory GCASP11 and 12 sequences are absent from *Cs* CAD1 and CAD5. None of the 12 Nara gc5 core GCASP paralogs are located next to another GCASP paralog, although 7 were found within 2 genes of another secreted protein-encoding gene (Supplementary Table S4).

To determine if GCASP clustering is conserved in other *Colletotrichum* spp. apart from *Cf* Nara gc5, GCASP-encoding genomic loci were investigated in other PacBio-sequenced genomes. This revealed that orthologs of GCASPs 1-9, except for GCASP3, are also located in syntenic gene clusters in *Ca* Cg56 and *Cs* Cg363 (Fig. 5A, Supplementary Fig. S10B). In *Cs* Cg363, three clusters, including a pair of tandemly duplicated clusters, were identified (Supplementary Fig. S10B). Further, in *Ca* Cg56, a GCASP-encoding cluster is present in the subtelomeric region of 56_c05 and appears to have been acquired in a lineage-independent manner since it is absent from homologous regions in *Cf* Nara gc5, Cf413 and *Cs* Cg363 (Fig. 5A). Analysis of read mapping depths indicates that, as in *Cf* Nara gc5, GCASP genes are present in multiple copies in pathogenic strains of *Cf* (S1 and S4), *Cs* (Cg363, CAD1, and CAD5), and *Ca* (Cg56) (Fig. 5B) and suggests that the number of clusters identified in *Cs* Cg363 and *Ca* Cg56 may be underestimated, possibly due to the difficulty of assembling these regions.

**Figure 5.**
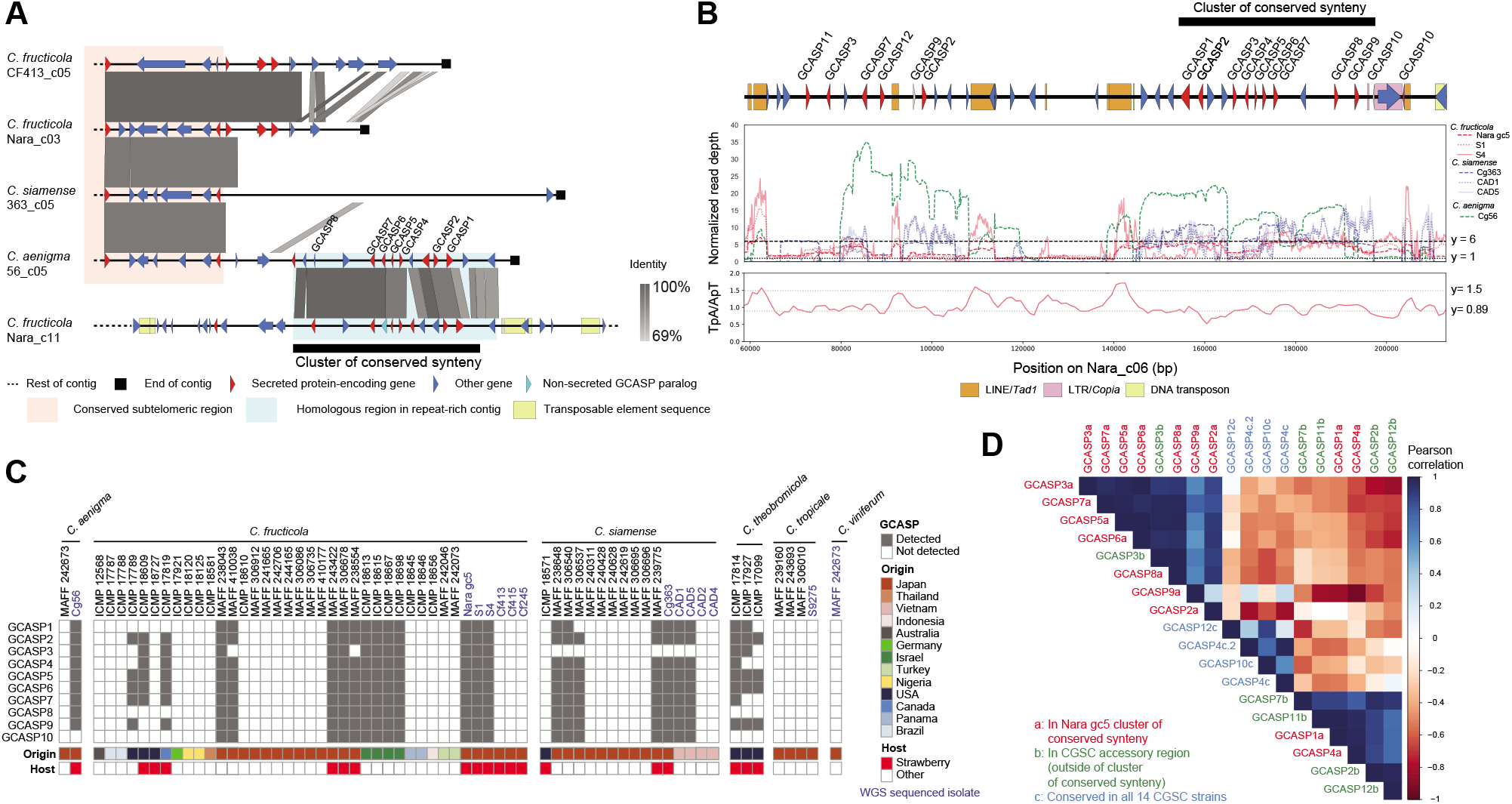
GCASP-encoding clusters in *Colletotrichum gloeosporioides* species complex isolates. (A) Conservation of a GCASP-encoding cluster at the subtelomeric region of *C. aenigma* Cg56. The absence of this cluster from subtelomeric regions which share synteny with this region in *C. fructicola* Nara gc5, Cf413, and *C. siamense* Cg363 is shown. (B) Normalized read mapping depths of a representative GCASP-encoding region in Nara gc5 showing the presence of related sequences in other isolates. TpA/ApT ratios are an indication of repeat-induced point mutation at this locus. (C) Conservation of sequences GCASP homologs in different CGSC species from a broad geographical distribution based on PCR. (D) Pearson correlation scores between expression profiles of different GCASP sequences in Nara gc5 conidia and during infection of *F. vesca* at 1, 3, and 6 days post-inoculation (dpi). Sequences associated with accessory clusters of conserved synteny show a tendency for correlated gene expression.

The conservation of GCASP sequences associated with clusters of conserved synteny (GCASPs 1-10) was further assessed by PCR in an additional 51 CGSC isolates from different geographical locations and hosts (Fig. 5C, Supplementary Fig. S10). In total, 26 isolates were identified with at least four GCASP-related sequences. This analysis revealed that these sequences are conserved in different CGSC species, namely *Cf* (16 out of 40), *Cs* (7 out of 16)*, Ca* (1 out of 2) and *C. theobromicola* (3 out of 3), with isolates in *Cf, Cs* and *Ca* showing presence/absence polymorphisms. Further, GCASP genes were detected in isolates from different hosts, namely, strawberry, apple, *Limonium* spp. and cassava. Isolates identified with GCASP1-10 sequences originated from diverse geographic locations, such as, Canada, the United States, Israel, Japan and Vietnam (Figs. 5c).

### GCASP genes are upregulated *in planta* and GCASP paralogs associated with clusters of conserved synteny tend to show correlated gene expression

To assess if GCASPs have a role in infection, we examined their expression in three different species, *Cf* Nara gc5, *Ca* Cg56 and *Cs* Cg363. Primers for quantitative PCR were designed for selected *Cf* Nara gc5 sequences (Supplementary Figs. S11-12). These results indicate that GCASP2-8 sequences are expressed *in planta* in *Cf* Nara gc5, suggesting a potential role for these genes in infection. In addition, we calculated pairwise correlations for all tested GCASP sequences within each strain (Fig. 5d, Supplementary Fig. S13). This revealed that the expression of *Cf* Nara gc5 GCASP paralogs in clusters of conserved synteny tended to show a more positive correlation to other GCASPs within clusters of conserved synteny rather than to paralogs located outside these regions (Fig. 5d). Interestingly, similar tendencies for correlated gene expression of cluster-associated GCASPs especially GCASPs 2, 5-7 were also shown in *Cs* Cg363 and *Ca* Cg56 (Fig. 5d, Supplementary Fig. S13).

## Discussion

The compartmentalization of fungal genomes into conserved and flexible regions is thought to allow the conservation of core, housekeeping genes, while allowing other genes to evolve rapidly, maintaining their pathogenicity, and avoiding host recognition. Studies have shown that the genomes of *Colletotrichum* spp. such as *C. gloeosporioides* (He et al. 1998) and *C. higginsianum* (Plaumann et al. 2018), include core, chromosomes, and repeat-rich accessory minichromosomes that can be gained or lost with little effect to vegetative growth. Despite the demonstration of minichromosome transfer between vegetative incompatible isolates more than 20 years ago (He et al. 1998), only recently has pathogenicity been linked to the presence of specific minichromosomes in certain strains (Plaumann et al. 2018; Bhadauria et al. 2019). In these recent studies, the minichromosome sequences were shown to be virulence determinants on host plants. However, as both types of chromosomes exist in the same nucleus, TE-dense accessory chromosomes may also play an additional role, affecting “core” chromosomes through the exchange of genes and promoting recombination.

Our study shows evidence for such a role of repeat-rich minichromosomes in the CGSC. The *Cf* Nara gc5 chromosome, Nara_c11, which is conserved in a subset of closely related *Cf* strains (Nara gc5, S1 and S4), has traits of both “core” and “accessory” chromosomes. Based on whole genome alignments with highly contiguous genome assemblies from other CGSC strains generated in this study, we propose that this chromosome originated from the recombination of a conserved “core” chromosome with a non-conserved accessory chromosome. Furthermore, our results strongly support the exchange of genes between subtelomeric regions of “core” chromosomes and the repeat-rich compartment of Nara_c11, resulting in the expansion of a group of candidate effector genes that are organized in clusters. The role of subtelomeres as compartments for effector duplication and diversification has been reported in *Magnaporthe oryzae*, where copies of the effector gene *AvrPita* are present in subtelomeric regions and accessory chromosomes(Orbach et al. 2000; Chuma et al. 2011). It is thought that the presence of these genes at these loci contribute to the observed frequent loss and mutation of *AvrPita*, which is recognized by hosts expressing the *Pi-ta R* gene (Orbach et al. 2000). Intriguingly, the presence of related effector candidate gene clusters in different subtelomeric loci in *Cf* and *Ca* indicates that these have been gained and/or lost independently in different lineages, despite likely sharing a common origin. This is reminiscent of recently described subregions of *Verticillium dahliae* and *Verticillium tricorpus* effector-coding “lineage-specific regions” which have absence/presence polymorphisms in different strains of the same species, but are conserved between different *Verticillium* species (Depotter et al. 2019). As in the case of the CGSC accessory regions, these regions show high sequence similarity between different *Verticillium* species and were proposed to originate prior to speciation. However, unlike the CGSC accessory regions, they were not found to be enriched in subtelomeric regions or to be associated with multicopy effector clusters.

Effector gene clusters have been observed in other plant pathogenic fungi, such as *Ustilago maydis* (Kämper et al. 2006). Recently, the PWL2 and BAS1 effectors that are only present in subtelomeres of different core chromosomes in the rice pathogen *M. oryzae* (MoO), were found to exist side-by-side only in dispensable, minichromosomes of wheat-adapted *M. oryzae* (MoT) indicating a potential role for minichromosomes as a compartment for accumulating pathogenicity-related sequences (Peng et al. 2019). Half of the GCASPs have paralogs that are also present in core genomic regions outside of accessory regions, supporting the enrichment of effector candidates from different loci in *Colletotrichum*.

While there are parallels between *M. oryzae* and the CGSC genomes, the minichromosomes of *M. oryzae* do not experience significant amounts of RIP mutations (Peng et al. 2019), unlike the dispensable region of Nara_c11 and the potential minichromosome, Nara_c12. In *Leptosphaeria maculans*, RIP suppresses expression of repeat-associated genes, although neighboring genes can be expressed (Rouxel et al. 2011). It is tempting to speculate that in the CGSC, the presence of RIP in regions flanking accessory GCASP loci suppresses the expression of genes from any single cluster, driving the need for multiple copies for increased pathogenicity. Although other genetic effects cannot be excluded, such a dosage effect may indicate why pathogenic CGSC strains encode multiple copies of these genes. The expression analysis also indicates that GCASP paralogs in clusters of conserved synteny tend to show similar expression dynamics compared to non-clustered paralogs. This is in line with the hypothesis that organization of these genes in clusters may be partially driven by the advantage of co-regulation, while maintaining them in a repeat-rich environment. However, this hypothesis needs to be further investigated.

The large-scale rearrangements, which may have generated Nara_c11, not only produce genetic diversity, such as chimeric gene sequences, but also affect the 3D organization of the genome, with potential effects on gene accessibility and expression (Spielmann et al. 2018). Chromosomal rearrangements have been observed in other plant pathogenic fungi such as *V. dahliae* and also *C. higginsianum* (Jonge et al. 2013; Tsushima et al. 2019). However, *V. dahliae* and *C. higginsianum* are asexual pathogens, whereas CGSC members have known sexual morphs (Weir et al. 2012). Indeed, it is noteworthy that *Cs* Cg363 and *Ca* Cg56 are highly similar to *Cf* Cf413 in terms of their genome organization despite belonging to different lineages. Therefore, the rearrangements observed may result in the reproductive isolation of the *Cf* Nara gc5 lineage from other *Cf* strains. Additionally, the fact that RIP occurs during the sexual cycle, may also cause increased TE activity in Nara gc5, although this needs to be investigated. It is noted that extensive rearrangements have also been observed during vegetative growth of the sexual fungal pathogen *Zymoseptoria tritici* (Möller et al. 2018).

Taken together, our results highlight the importance of accessory sequences in subtelomeric and repeat-rich chromosomes in increasing genome plasticity in *Colletotrichum* and add an additional dimension to the role of these sequences in affecting the genome evolution in this group of important plant pathogens.

## Materials and methods

### Fungal cultures

Details of all fungal strains used can be found in Supplementary Table S5. For genomic DNA extraction, fungi were cultured in potato dextrose broth (BD Biosciences, Franklin Lakes, NJ, USA) at 24°C in the dark for 2 days. For production of conidia, fungi were incubated on Mathur’s media or potato dextrose agar (Nissui Pharmaceutical Co. Ltd., Tokyo, Japan) at 24°C for 12 h under black-light blue fluorescent bulb light/12 h dark conditions for ten days. Conidia were released in autoclaved distilled water, filtered through a 100 μm cell strainer (BD Biosciences) and pelleted by centrifugation at 3300 × g for 5 min before resuspension in autoclaved distilled water to the final desired concentration.

### Plant infections

For infection of *F*. × *ananassa* var. Sachinoka, leaves were detached from strawberry plantlets grown under long day conditions (16 h light/8 h dark) at 25°C and inoculated with 5 μL droplets of 5 × 10^5^ conidia/mL conidial suspensions of each strain. Infected leaves were maintained at a 100 % humidity under 12 h light/12 h dark conditions at 22°C. Pathogenicity was assessed from 5 days post-inoculation (dpi). For infection of *F. vesca*, 5- to 6-week-old plants grown under long day conditions (16 h light/8 h dark) at 25°C were spray-inoculated with conidial suspensions at the specified concentrations. Plants were kept at a 100 % humidity under long day conditions prior to imaging. For drop inoculations of *F. vesca* plants, the fifth and sixth leaves from 5- to 6-week-old plants were detached and inoculated with 5 μL of 5 × 10^5^ conidia/ml conidial suspensions. Lesions were assessed at 7 dpi.

### Genome sequencing, assembly and annotation

Genomic DNA was isolated using CTAB and 100/G genomic tips (QIAgen, Hilden, Germany) as described in the 1000 Fungal genomes project (http://1000.fungalgenomes.org). Details on library preparation, sequencing and assembly are in Supplementary Table S6. Annotations for *Cf* strains Nara gc5, S1 and Cf413 were generated using the BRAKER1 pipeline using hints from RNAseq reads, mapped to each genome using hisat2 with –max-intronlen set to 1000. *Ca* Cg56 and *Cs* Cg363 were annotated using the MAKER2 pipeline using gmes parameters trained on *Cf* Nara gc5 and Augustus parameters trained on each genome using BUSCO v3 with sordariomycete conserved genes. Annotations were prepared for NCBI using the GAG (Geib et al. 2018) and Annie (Tate et al. 2014) annotation pipelines. Assemblies were assessed with quast v4.6.3 (Gurevich et al. 2013) using the default settings and BUSCO v3 (Simão et al. 2015) using pezizomycotina_odb9 gene models and the settings: “--long --fungal”. PFAM domains from Pfam release 32 (Aug 2018) were assigned to predicted proteins of *Cf* Nara gc5 and Cf413; *Ca* Cg56, *Cs* Cg363 and *C. higginsianum* by performing pfam_scan (v1.6) using the cut_ga gathering threshold and -as settings. The localizations of annotated fungal proteins were predicted using DeepLoc v1 (Almagro Armenteros et al. 2017). In addition, EffectorP v1 and 2 (Sperschneider et al. 2016; Sperschneider et al. 2018) were run on all predicted extracellular proteins. Predicted proteins were assigned as CAzymes using dbCAN (Yin et al. 2012), as proteases using the MEROPS database (Rawlings et al. 2012) and as secondary metabolites using antiSMASH (Blin et al. 2013) and SMIPS (Wolf et al. 2016).

### Analysis of repeat elements

Repeats from *Cf* Nara gc5, *Cf* Cf413, *Cs* Cg363, *Ca* Cg56 and *C. higginsianum* IMI 349063 were predicted using RECON and RepeatScout via RepeatModeler v open-1.0.11 (Smit and Hubley 2008), TransposonPSI (http://transposonpsi.sourceforge.net/), LTR_retriever (Ou and Jiang 2018) and LTRPred (Benoit et al. 2019) (https://github.com/HajkD/LTRpred) as described in our project repository. Sequences that were longer than 400 bp from TransposonPSI, LTR_retriever and LTRPred were combined and used as queries for BLASTx against RepBase peptide v23.12 (Bao et al. 2015) sequences included with RepeatMasker (v 1.322). Further, sequences were used as queries for BLASTn against each fungal genome. Only sequences with more than five hits (BLASTn *E-*value cutoff of 1E-15) and/or with hit to a RepBase peptide (BLASTx *E-*value cutoff=1E-5) were retained for further analysis. Sequences from all sources were combined using vsearch, by using an 80 % identity as the cutoff threshold. Consensus sequences were classified using RepeatClassifier (from RepeatModeler) and the genome was masked using the custom repeat library using RepeatMasker. The “one code to find them all” was used to reconstruct transposable elements (Bailly-Bechet et al. 2014). Custom scripts were used to determine the location of the telomeric repeat TTAGGG and to calculate TpA/ApT dinucleotide frequencies over 3 kb windows.

### Read coverage analysis

Bowtie2 v2.3.4.1 (Langmead and Salzberg 2012) was used to map Trim Galore quality-trimmed paired-end reads from 500 bp insert libraries to assemblies using the settings: “--end-to-end –fr – very-sensitive -X 700 -I 400”. Duplicate reads were removed using the samtools v1.8 (Li et al. 2009) fixmate and markdup with default settings. The number of mapped reads per 10 kb was obtained using the bedtools v2.29.2 (Quinlan and Hall 2010) coverage tool and normalized over genome-wide median read depths. To estimate the number of GCASP-associated clusters, reads were mapped to Nara_c06, which has only a single copy of the GCASP cluster, as described and read depths were calculated using bedtools coverage. Read depths were normalized over the median read depths of Nara_c06 from position 2,000,000 to 4,000,000, since this region is present as a single copy in the genome.

### Enrichment analysis

Enrichment scores were calculated by performing simulations as previously described (Nègre et al. 2010). In brief, locations of features of interest were randomly permuted within each chromosome of origin using the pybedtools randomstats function with the “-chrom” setting. A total of 10,000 simulations were assessed for proportions of overlaps with each set of features of interest. Enrichment scores were calculated by normalizing the actual proportions of overlaps between the two sets of features by the median of the simulated, randomized dataset. Empirical P-values were obtained by determining the fraction of simulated overlaps that are greater than the observed overlap. Telomeric repeat-rich regions were defined as 100 kb windows with TTAGGG densities greater than 95 % of the genome.

### Identification of large-scale structural genomic rearrangements

Whole genome alignments were conducted to identify large scale structural genomic changes between each genome assembly using nucmer from the mummer suite of programs with the “maxmatch” setting (Delcher et al. 2003). This was followed by filtering sequence lengths in the reference sequence as defined in the text. Circos (v0.69.6) and mummerplot (v3.5) were used to visualize the genomic rearrangements. Regions of synteny were identified using SynChro (Drillon et al., 2014) from the CHROnicle package with a delta of 3 as previously described (Shi-Kunne et al. 2018). Plots of specific syntenic regions were obtained using EasyFig v2.2.2 (Sullivan et al. 2011). In addition, the mapping of reads at identified synteny breaks of greater than 100 kb between *Cf* Nara gc5 and Cf413 generated from bowtie2 mappings described above were visually examined using CLC GenomicsWorkbench v12 (QIAGEN, Hilden, Germany). For strains with available PacBio reads, long reads were mapped to the *Cf* Nara gc5 genome using ngmlr v0.2.7 (Sedlazeck et al. 2018) by using default settings.

### Orthogroup and phylogenetic analyses

Orthofinder2 (Emms and Kelly 2019) was used to identify orthogroups in the predicted proteins of the 33 fungal species tested (Supplementary Table S4) by using the default settings. For *F. verticillioides, F. oxysporum*, *V. dahliae* and *N. crassa*, only T0 transcript models from each locus were included. OrthoFinder2 was also used to generate the rooted species tree of the 33 fungal species. Maximum likelihood phylogenetic analyses were carried out by RAxML-ng (Kozlov et al. 2019) on GCASP orthogroups after determining the best models for substitution using modeltest-ng (Darriba et al. 2020). For each GCASP orthogroup, the best scoring tree of 500 parsimonious and 500 random trees was used with support values from 1,000 bootstrap replicates. For maximum likelihood phylogeny of all *Colletotrichum* members, whole genome SNPs were identified from nucmer whole genome alignments by PhaME (Shakya et al. 2020). Maximum likelihood phylogenetic analysis was carried out using RAxML-ng on the 199,953 SNP positions identified with the GTR+G4 model of substitution, which was determined as the best model according to modeltest-ng (Darriba et al. 2020).

### Expression analysis

RNA from detached *F*. × *ananassa* var. Sachinoka leaves inoculated with 5 × 10^5^ conidia/mL at 1 dpi, 3 dpi and 6 dpi, as well as 2 dpi roots of intact *F*. × *ananassa* var. Sachinoka plantlets grown in vermiculite under 12 h light/12 h dark conditions at 22°C were isolated using the improved 3% CTAB3 method (Yu et al. 2012) followed by treatment with RNase-free DNase (QIAGEN, Venlo, Netherlands) and clean up with RNeasy spin columns (QIAGEN, Venlo, Netherlands). RNA from *in vitro* conidia and hyphae harvested 3 and 7 days post-inoculation of potato dextrose broth grown in the dark at 24°C were extracted with RNeasy spin columns (QIAGEN, Venlo, Netherlands). Illumina TruSeq RNAseq libraries were prepared according to the manufacturer’s instructions and sequenced on an Illumina HiSeq 2500 sequencer (50 bp single reads). Reads from 3 biological replicates per sample were mapped to the *Cf* Nara gc5 reference genome using STAR (Dobin et al. 2013)(v2.6.0a) with the setting: “--alignIntronMax 1000”. Read counts were obtained using Rsubread (Liao et al. 2019) (v1.32.2) with the “primaryOnly=TRUE” and “strandSpecific=2” settings. Genes were considered to have evidence for expression if they passed the EdgeR (Robinson et al. 2010) filterByExpr function. Using the EdgeR package, filtered read counts were TMM normalized and the glmQLFtest was used to identify *in planta* up-regulated genes as genes that were differentially up-regulated (FDR<0.05) at 1, 3 or 6 dpi on detached leaves or 2 dpi on roots relative to 3 day *in vitro* grown hyphae.

For quantitative RT-PCR, RNA from conidia and 1, 3 and 6 dpi-detached, spray-inoculated *F. vesca* leaves with 5 × 10^5^ conidia/mL conidial suspensions of *Cf* Nara gc5, *Cs* Cg363 or *Ca* Cg56 were isolated as described for *F*. × *ananassa*. cDNA was generated using the ReverTra Ace reverse transcriptase (Toyobo Co., Ltd., Osaka, Japan) using random primers for priming. Transcripts were detected using THUNDERBIRD SYBR qPCR mix (Toyobo Co., Ltd., Kita-ku, Osaka, Japan) on a MX3000P Real-Time qPCR System (Stratagene, Santa Clara, California, USA) with primers specified in Supplementary Table S7. Transcript levels were quantified according to the standard curve method and genes were normalized to the expression of *CfEF*. Relative expression values were scaled using the MinMaxScaler from Sklearn (v0.20.3) in a gene-wise manner. After excluding genes with no change in expression patterns and/or genes that were not detected at any of the tested stages, Pearson correlation scores were calculated in a pairwise manner using the R stats package (v3.5.3) and visualized using the corrplot package (v0.84).

### PCR amplification of GCASP sequences from non-sequenced CGSC isolates

PCR was carried out to confirm the conservation of related GCASP sequences in DNA isolated from the additional 51 strains. PCR reactions were carried out according to manufacturer’s instructions using either ExTaq polymerase (TaKaRa Bio, Tokyo, Japan) or PCR master mix (Promega, Wisconsin, USA). For ExTaq polymerase reactions, 0.625 U of ExTaq polymerase (TaKaRa Bio, Tokyo, Japan), 0.2 μM primers (Supplementary Table S7), 0.16 mM dNTPs and 0.5-10 ng of purified genomic DNA. Tubes were heated at 94°C for 3 min, followed by 30 cycles of denaturation at 94°C for 30 s, annealing at 55-60°C for 30 s and extension at 72°C for 1 min, followed by a final extension at 72°C for 5 min. For PCR master mix reactions, 10 μl of 2×PCR master mix was mixed with 0.5 μM primers and 0.5-10 ng of genomic DNA. Tubes were heated at 95°C for 5 min, followed by 35 cycles of denaturation at 95°C for 30 s, annealing at 55-60°C for 30 s and extension at 72°C for 1 min, followed by a final extension at 72°C for 5 min. For each amplification, genomic DNA from *Cf* Nara gc5 and Cf413 were included as positive and negative controls, respectively.

## Data availability

Assemblies and raw reads are deposited in GenBank. More detailed descriptions and code for running analyses are available in: https://github.com/pamgan/colletotrichum_genome

## Acknowledgements

Newly generated assemblies (Accessions: QLYQ00000000, QPMV00000000, ANPB02000000, QPMY00000000, QPMX00000000, QPMW00000000, QPNA00000000, QPMU00000000, QPMT00000000, QPMS00000000, QPMR00000000, QPMP00000000, QPNB00000000, QPMQ00000000) and raw reads (SRA accession PRJNA476648) are deposited in GenBank. We would like to thank Dr. Takeshi Suzuki and Dr. Nanako Nakata for providing strains S1, S4, Cf413, Cf415, Cf245, Cg363 and Cg56 and Dr. Anuphon Laohavisit for critical reading of the manuscript. This work was supported by the Grant-in-Aid for Scientific Research (KAKENHI) (17H06172 and 15H05959 to K.S., 18H02204 to Y.T., 19K15846 to P.G.), the Science and Technology Research Promotion Program for the Agriculture, Forestry, Fisheries and Food industry to Y.N., Y.T. and K.S. and JSPS DC2 fellowship to A.T.

## Supplementary Material

**Supplementary Fig. S1.** Infection of *Fragaria vesca* by strains from the *Colletotrichum gloeosporioides* species complex. At 3 days post-inoculation (dpi) (A, B, C) appressoria (filled arrowheads) have penetrated leaf epidermal cells to form penetration pegs (A) infection vesicles (B, C) and intracellular hyphae (C). At 5 dpi (D) intercellular secondary hyphae proliferate. (E) Symptoms of infection of the 14 sequenced strains on *F. vesca* at 7 dpi. (F) Spray-inoculated *F. vesca* plants at 4 dpi. Scale bars for A-C: 20 μm; D: 50 μm V: infection vesicles

**Supplementary Fig. S2.** Infection of *Fragaria* × *ananassa* var. Sachinoka by the 14 sequenced isolates from the *Colletotrichum gloeosporioides* species complex at 7 days postinoculation (dpi) (A). (B) *In planta* infection by *C. fructicola* Nara gc5 and Cf413. Conidia of both strains develop appressoria by 1 dpi. At 3 dpi, intracellular hyphae are observed in Nara gc5 infections (filled arrowheads), but not in Cf413. At 6 dpi, secondary hyphae proliferate in Nara gc5 infections but not in Cf413, although conidia remain metabolically active and able to express GFP.

**Supplementary Fig. S3.** Relationship of strains with other known sequenced isolates. (A) Maximum likelihood phylogeny of *Colletotrichum* fungi based on genome-wide SNPs identified by PhaMe by nucmer alignment. 199,953 SNPs were concatenated, and the phylogeny was estimated based on the GTR+G4 model using raxml-ng. The most likely tree out of 100 random and 100 parsimony-based trees is shown with node bootstrap support values out of 100 replicates. Trees converged after 50 bootstraps. (B) Nucmer global alignment of Nara_c13 shows homology with KX034082.1 which encodes the full *C. fructicola* mitochondrial genome (KX034082; Liang et al. 2017). Red lines indicate forward matches while blue lines indicate reverse matches.

**Supplementary Fig. S4.** (A) Heatmaps showing hierarchical clustering of genes encoding extracellular carbohydrate active enzymes (CAzymes) in 33 fungal genomes by gene copy number. CAzymes were defined as proteins that were identified by at least two tools (HMMER, DIAMOND and/or HotPep) in dbCAN (Yin et al. 2012). (B) Heatmap showing hierarchical clustering of copy numbers of genes encoding extracellular proteases and protease inhibitors in 33 fungal genomes. Proteases and protease inhibitors were identified with BLASTp against protease and protease inhibitor domains from the MEROPS database. The localizations of proteins were determined by analysis with DeepLoc. (C) Bar plot showing secondary metabolite (SM) genes identified by analysis with SMIPS. Strain-specific expansion of acyltransferase (trans-AT) type SM genes was observed in *Colletotrichum fructicola* Nara gc5. Numbers of genes in each category are given. The key to abbreviations is given in Supplementary Table S5. Names of strains from the Colletotrichum gloeosporioides species complex are in red.

**Supplementary Fig. S5** Mapping of reads around genomic regions representing 14 synteny breaks (BP01-14) of greater than 100 kb between the *Colletotrichum fructicola (Cf)* Nara gc5 and the Cf413 genome assemblies identified by nucmer (maxmatch, 10 kb cutoff). Tracks depict 500 bp insert-sized paired-end read mappings by bowtie2, except for Nara gc5, S1, Cf413, Cg363 and Cg56, where ngmlr was used to align PacBio reads. Schematics above the mapping tracks show the point of synteny breaks relative to the whole contig. Read mapping indicates that genomic regions around Nara gc5 BP11, 13-14 are also present in *Cf* S1 and S4 and that genomic regions around Nara gc5 BP1 and 12 are also present in *Cf* S4.

**Supplementary Fig. S6.** Duplications in *Colletotrichum fructicola* Cf413 occur mostly between the ends of contigs (A) except for a single mid-contig segmental duplication that shows (B) synteny with the end of 413_c01. (C) Median normalized read mapping depths of contig 413_c09 shows that the region downstream of the mid-contig segmental duplication is duplicated in the Cf413 genome. Mean read depths per 10 kb are shown with 1 kb sliding window intervals.

**Supplementary Fig. S7.** Circos plot of homologous regions between 413_c05 and c09 and Nara_c03 and c10.Tracks from outermost to inner: 1: presence of reciprocal best blast hit in 413_c05 (blue) and 413_c08 (pink), 2: repeat density, 3: secondary metabolite biosynthesis gene clusters. Ribbons show nucmer aligned regions indicating the presence of segmental duplication where at the point of potential chromosomal rearrangement. Ticks represent 0.5 Mb.

**Supplementary Fig.S8.** Repeats in *Colletotrichum* species. Linear regression modeling of the relationship between the percentage of genome covered by repeats and the number of genes (A) and genome length (B). (C) Repeat content in *C. fructicola* Cf413 and *C. siamense* Cg363.

**Supplementary Fig. S9.** Features of all contigs in *Colletotrichum fructicola* Nara gc5 >100 kb in length. Number of reads mapping/10kb were normalized relative to whole genome medians. *In planta* upregulated: number of genes/10 kb that are significantly up-regulated at either 1, 3 or 6 days post-inoculation (dpi) during infection of *Fragaria* × *ananassa* leaves or 2 dpi *F*. × *ananassa* root tissue compared to 3 day-old *in vitro* hyphae. Spaces between ticks represent 1 Mb. Effector: number of genes predicted to encode effectors/10 kb.

**Supplementary Fig. S10.** *Colletotrichum gloeosporioides* species complex accessory regions. (A) Synteny between six GCASP-encoding gene clusters in *C. fructicola* Nara gc5. (B) Conservation of gene order in three syntenic clusters of conserved synteny encoding GCASP orthologs in *C. siamense* Cg363. Only hits of 500 bp or more and with less than 0.0001 E-value are visualized. (C) and (D) PCR to amplify GCASP-related sequences in 51 additional *C. gloeosporioides* species complex isolates, which is shown in Fig. 5C. A 100 bp or 1 kb ladder was used to provide size estimates. See Supplementary Table S5 for details of the strains used.

**Supplementary Fig. S11.** Maximum likelihood phylogenies of GCASP homologs with paralogs outside of accessory clusters of conserved synteny in *Colletotrichum fructicola* Nara gc5. GCASP1, 5 and 8 are not included as these sequences are only present in accessory clusters of conserved synteny. Values at nodes are percentages of support of 1000 bootstrap replicates. Purple labels: sequences used to design primers used for qPCR analysis (Supplementary Table S7 and Supplementary Fig. S12). The branch length of CSAL KXH65162.1 in GCASP11 is shortened 10-fold.

**Supplementary Fig. S12.** Quantitative PCR of selected GCASP homologs in *Colletotrichum fructicola (Cf)* Nara gc5, *C. aenigma* Cg56 and *C. siamense* Cg363 designed based on *Cf* Nara gc5 sequences in (A) accessory clusters of conserved synteny, (B) CGSC accessory regions and (C) regions that are conserved in all 14 sequenced CGSC strains. Expression levels relative to elongation factor 1 *(CfEF)* were assessed in conidia and in *F. vesca* infected leaves at 24, 72, and 144 hours post-inoculation (hpi). Data from 3-5 replicates per time point are shown with a boxplot showing the distributions of gene expression. Only sequences with at least one time point showing expression in any of the three strains are shown here.

**Supplementary Fig. S13.** Correlation plots to visualize the correlation of GCASP homolog genes expression in (A) *Colletotrichum aenigma* Cg56 and (B) *C. siamense* Cg363. Mean relative expression levels in each strain were scaled within each primer set and then pairwise correlation scores were calculated.

**Supplementary Table S1**. Features of contigs in the *Colletotrichum fructicola* Nara gc5 genome.

**Supplementary Table S2**. Enrichment of different features in CGSC accessory regions. Only PacBio-sequenced *Colletotrichum fructicola* and *C. siamense* strains were analyzed. **Supplementary Table S3.** *Colletotrichum gloeosporioides* species complex accessory secreted paralogs (GCASP) in *C. fructicola* Nara gc5. *: Proteins predicted to be effectors according to EffectorPv1 and/or v2. † Paralog that is not predicted to be secreted.

**Supplementary Table S4.** Features of genes in the *Colletotrichum fructicola* Nara gc5 genome. Extracellular sequences that are not predicted to be effectors are highlighted in green, and extracellular sequences that are also predicted to be effectors are highlighted in blue.

**Supplementary Table S5.** Fungal strains analyzed in this study.

**Supplementary Table S6.** Sequencing and assembly settings for *Colletotrichum*

*gloeosporioides* species complex members.

**Supplementary Table S7.** Sequences of primers used.

