## Supplementary Fig for "Subtelomeric regions and a repeat-rich chromosome harbor multicopy effector gene clusters with variable conservation in multiple plant pathogenic *Colletotrichum* species"

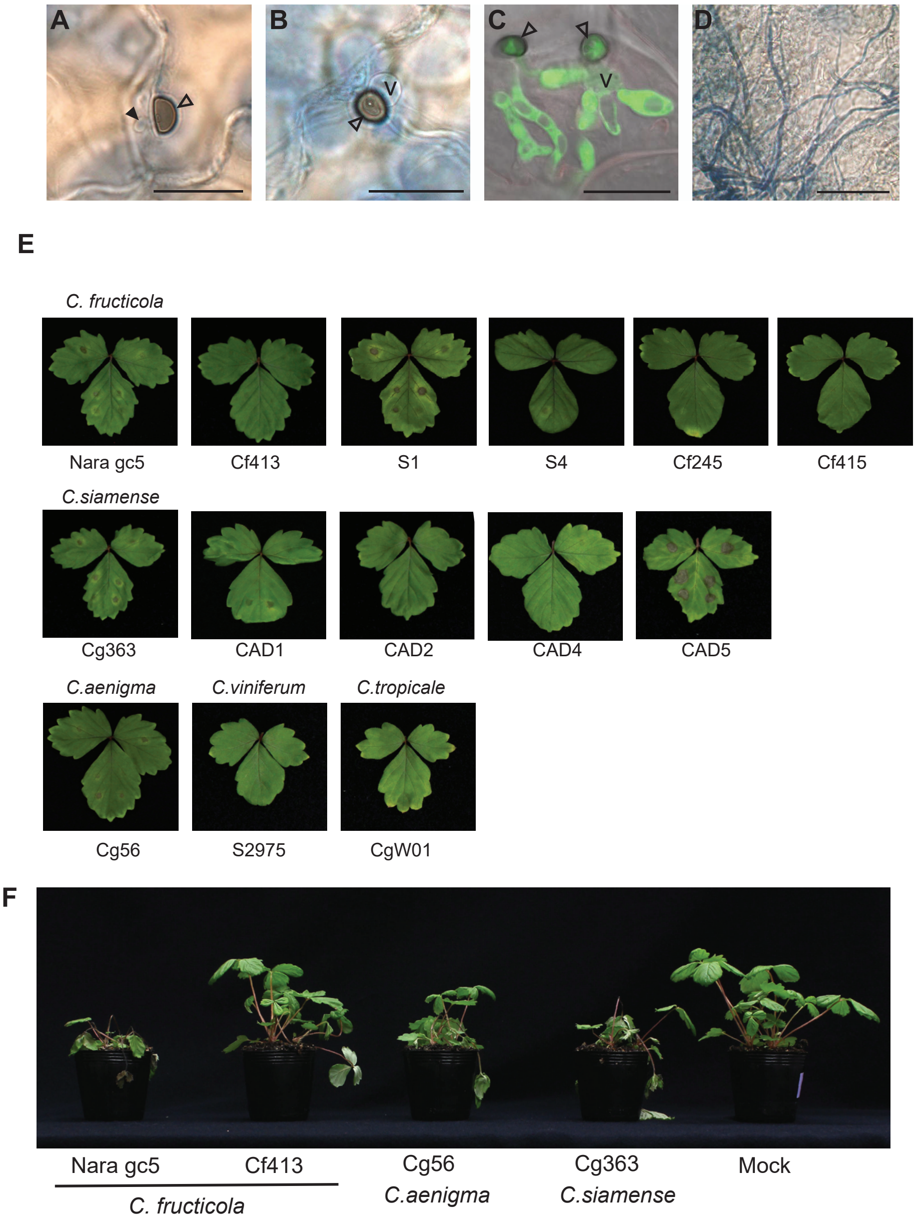


**Supplementary Fig. S1.** Infection of *Fragaria vesca* by strains from the *Colletotrichum gloeosporioides* species complex. At 3 days post-inoculation (dpi) (A, B, C) appressoria (filled arrowheads) have penetrated leaf epidermal cells to form penetration pegs (A) infection vesicles (B, C) and intracellular hyphae (C). At 5 dpi (D) intercellular secondary hyphae proliferate. (E) Symptoms of infection of the 14 sequenced strains on *F. vesca* at 7 dpi. (F) Spray-inoculated *F. vesca* plants at 4 dpi. Scale bars for A-C: 20 µm; D: 50 µm V: infection vesicles

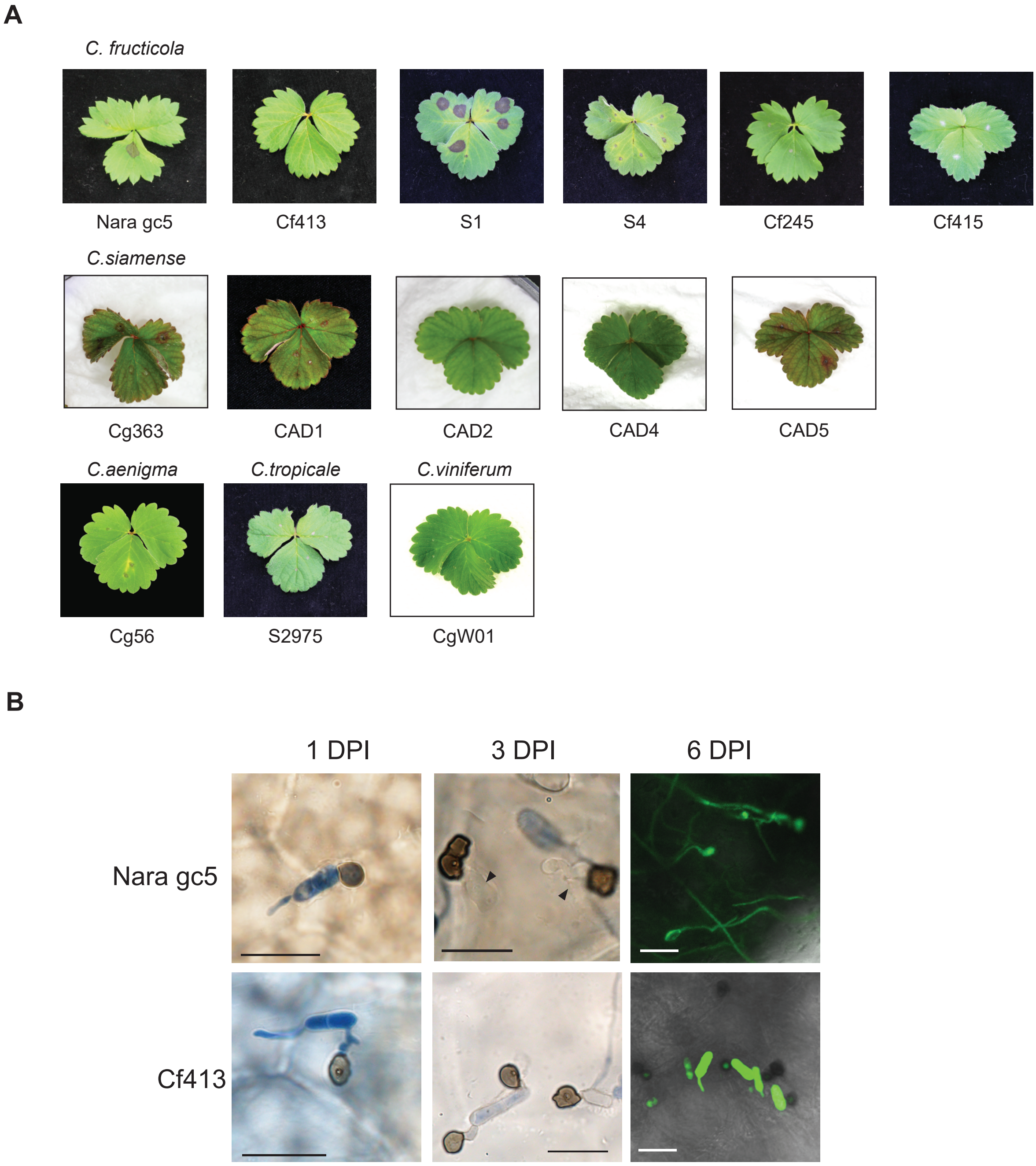


**Supplementary Fig. S2.** Infection of *Fragaria* × *ananassa* var. Sachinoka by the 14 sequenced isolates from the *Colletotrichum gloeosporioides* species complex at 7 days post-inoculation (dpi) (A). (B) In planta infection by *C. fructicola* Nara gc5 and Cf413. Conidia of both strains develop appressoria by 1 dpi. At 3 dpi, intracellular hyphae are observed in Nara gc5 infections (filled arrowheads), but not in Cf413. At 6 dpi, secondary hyphae proliferate in Nara gc5 infections but not in Cf413, although conidia remain metabolically active and able to express GFP.

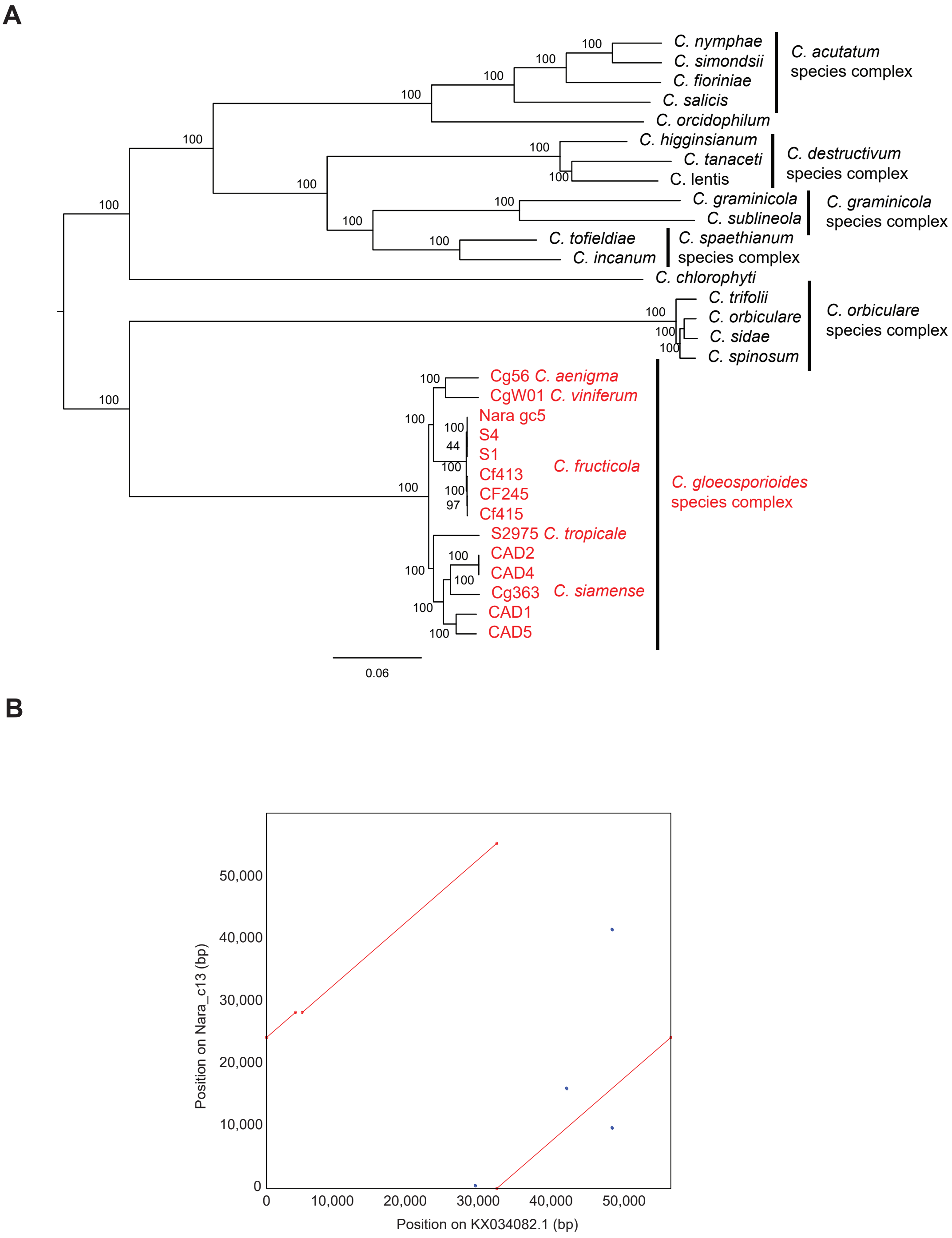


**Supplementary Fig. S3.** Relationship of strains with other known sequenced isolates. (A) Maximum likelihood phylogeny of *Colletotrichum* fungi based on genome-wide SNPs identified by PhaMe by nucmer alignment. 199,953 SNPs were concatenated, and the phylogeny was estimated based on the GTR+G4 model using raxml-ng. The most likely tree out of 100 random and 100 parsimony-based trees is shown with node bootstrap support values out of 100 replicates. Trees converged after 50 bootstraps. (B) Nucmer global alignment of Nara_c13 shows homology with KX034082.1 which encodes the full *C. fructicola* mitochondrial genome (KX034082; Liang et al. 2017). Red lines indicate forward matches while blue lines indicate reverse matches.

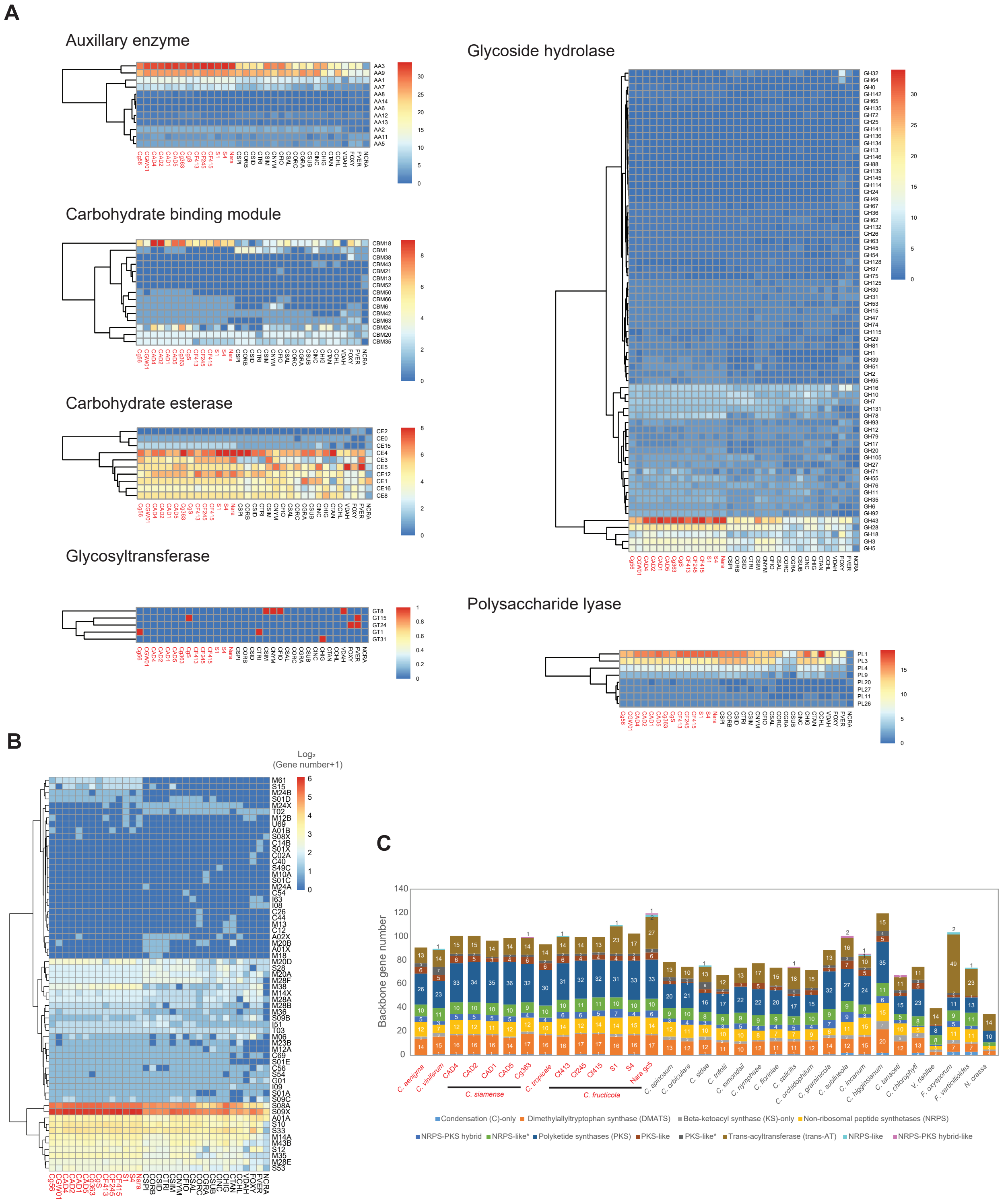


**Supplementary Fig. S4.** (A) Heatmaps showing hierarchical clustering of genes encoding extracellular carbohydrate active enzymes (CAzymes) in 33 fungal genomes by gene copy number. CAzymes were defined as proteins that were identified by at least two tools (HMMER, DIAMOND and/or HotPep) in dbCAN (Yin et al. 2012). (B) Heatmap showing hierarchical clustering of copy numbers of genes encoding extracellular proteases and protease inhibitors in 33 fungal genomes. Proteases and protease inhibitors were identified with BLASTp against protease and protease inhibitor domains from the MEROPS database. The localizations of proteins were determined by analysis with DeepLoc. (C) Bar plot showing secondary metabolite (SM) genes identified by analysis with SMIPS. Strain-specific expansion of acyltransferase (trans-AT) type SM genes was observed in *Colletotrichum fructicola* Nara gc5. Numbers of genes in each category are given. The key to abbreviations is given in Supplementary Table S5. Names of strains from the *Colletotrichum gloeosporioides* species complex are in red.

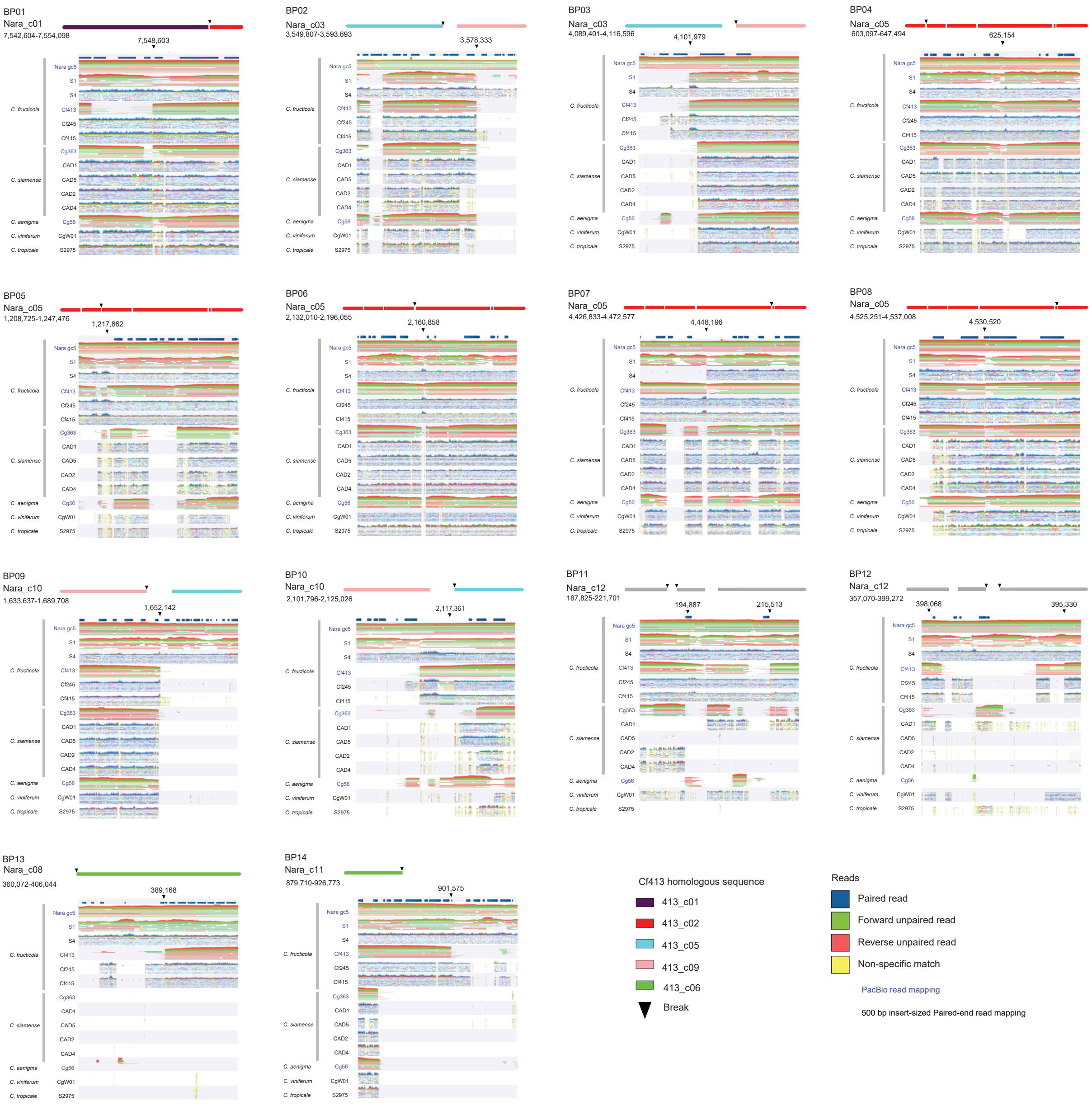


**Supplementary Fig. S5** Mapping of reads around genomic regions representing 14 synteny breaks (BP01-14) of greater than 100 kb between the *Colletotrichum fructicola* (*Cf*) Nara gc5 and the Cf413 genome assemblies identified by nucmer (maxmatch, 10 kb cutoff). Tracks depict 500 bp insert-sized paired-end read mappings by bowtie2, except for Nara gc5, S1, Cf413, Cg363 and Cg56, where ngmlr was used to align PacBio reads. Schematics above the mapping tracks show the point of synteny breaks relative to the whole contig. Read mapping indicates that genomic regions around Nara gc5 BP11, 13-14 are also present in *Cf* S1 and S4 and that genomic regions around Nara gc5 BP1 and 12 are also present in *Cf* S4

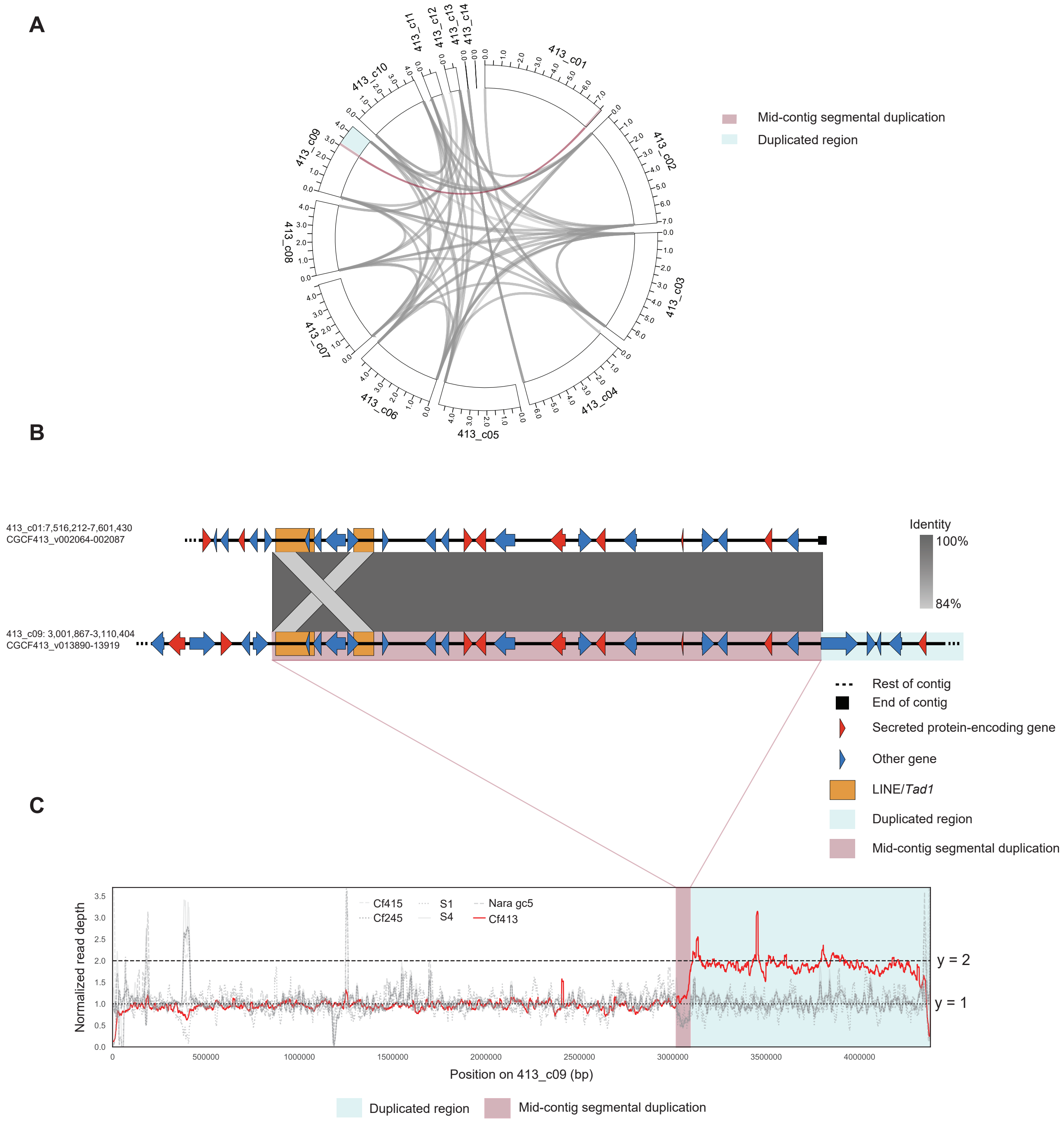


**Supplementary Fig. S6.** Duplications in *Colletotrichum fructicola* Cf413 occur mostly between the ends of contigs (A) except for a single mid-contig segmental duplication that shows (B) synteny with the end of 413_c01. (C) Median normalized read mapping depths of contig 413_c09 shows that the region downstream of the mid-contig segmental duplication is duplicated in the Cf413 genome. Mean read depths per 10 kb are shown with 1 kb sliding window intervals.

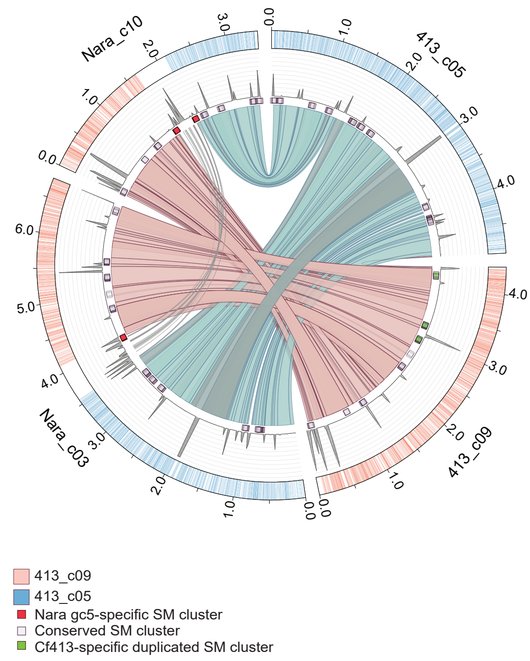


**Supplementary Fig. S7.** Circos plot of homologous regions between 413_c05 and c09 and Nara_c03 and c10.Tracks from outermost to inner: 1: presence of reciprocal best blast hit in 413_c05 (blue) and 413_c08 (pink), 2: repeat density, 3: secondary metabolite biosynthesis gene clusters. Ribbons show nucmer aligned regions indicating the presence of segmental duplication where at the point of potential chromosomal rearrangement. Ticks represent 0.5 Mb.

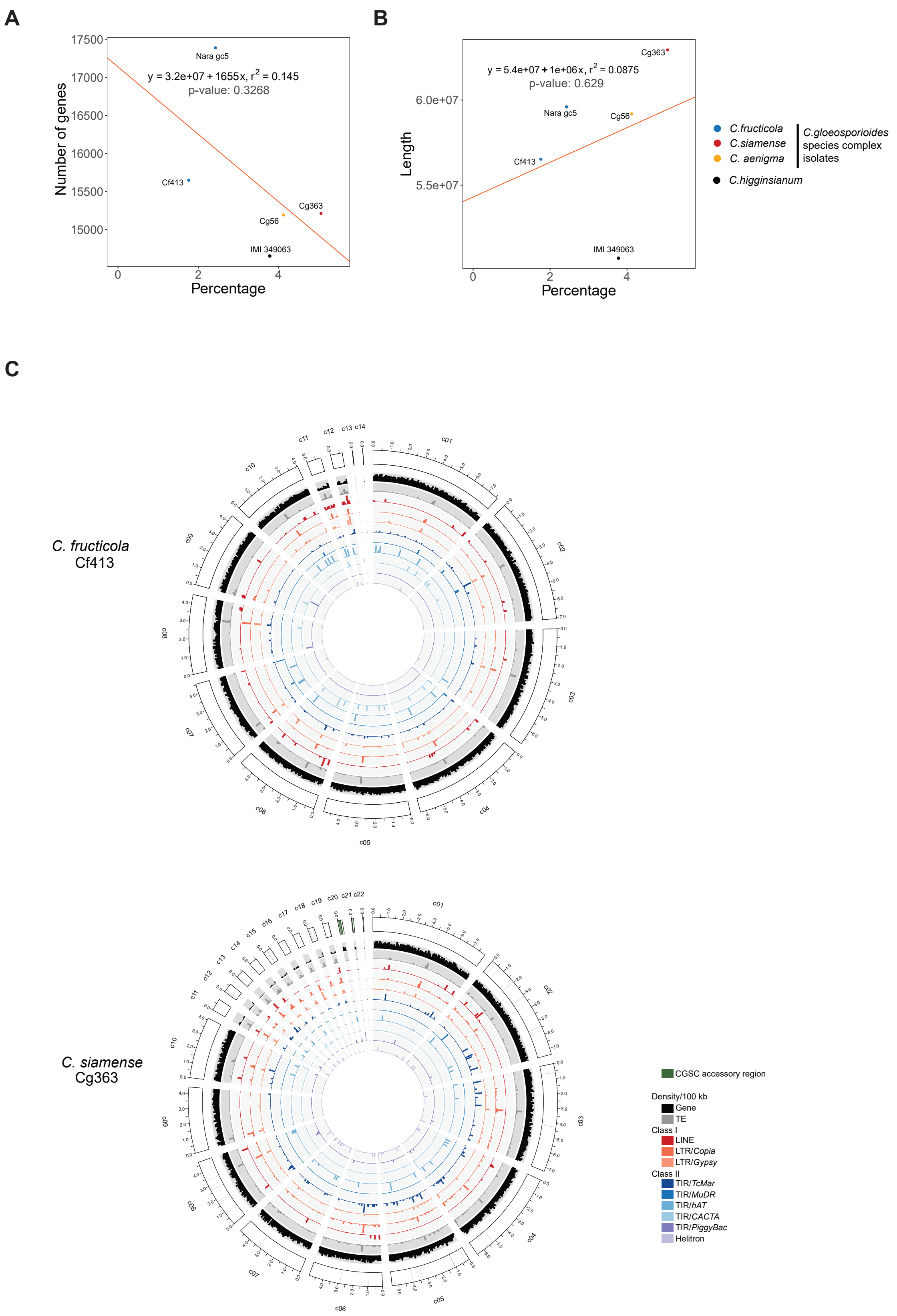


**Supplementary Fig.S8.** Repeats in *Colletotrichum* species. Linear regression modeling of the relationship between the percentage of genome covered by repeats and the number of genes (A) and genome length (B). (C) Repeat content in *C. fructicola* Cf413 and *C. siamense* Cg363.

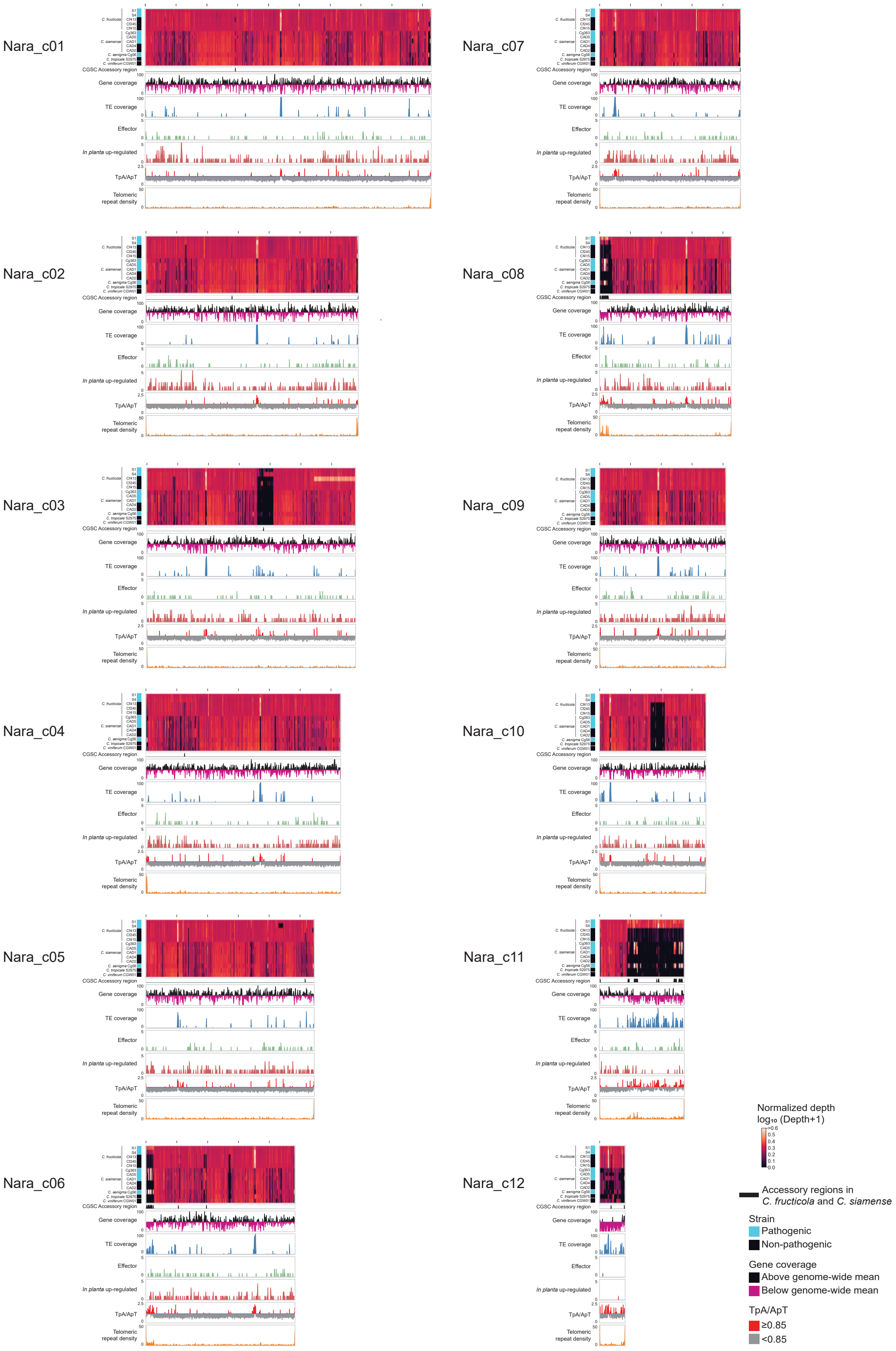


**Supplementary Fig. S9.** Features of all contigs in *Colletotrichum fructicola* Nara gc5 >100 kb in length. Number of reads mapping/10kb were normalized relative to whole genome medians. *In planta* upregulated: number of genes/10 kb that are significantly up-regulated at either 1, 3 or 6 days post-inoculation (dpi) during infection of *Fragaria* × *ananassa* leaves or 2 dpi *F.* × *ananassa* root tissue compared to 3 day-old *in vitro* hyphae. Spaces between ticks represent 1 Mb. Effector: number of genes predicted to encode effectors/10 kb.

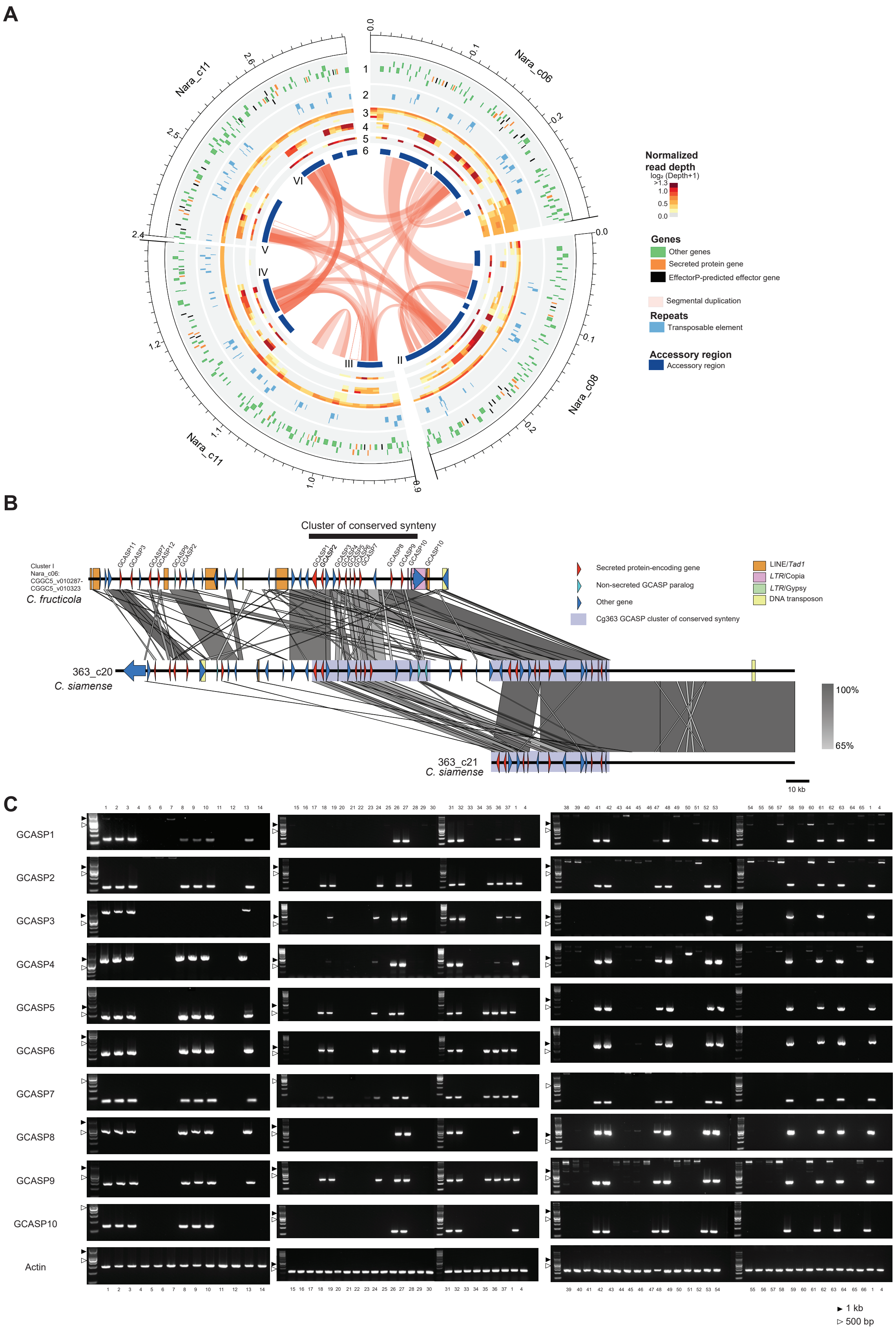


**Supplementary Fig. S10.** *Colletotrichum gloeosporioides* species complex accessory regions. (A) Synteny between six GCASP-encoding gene clusters in *C. fructicola* Nara gc5. (B) Conservation of gene order in three syntenic clusters of conserved synteny encoding GCASP orthologs in *C. siamense* Cg363. Only hits of 500 bp or more and with less than 0.0001 E-value are visualized. (C) and (D) PCR to amplify GCASP-related sequences in 51 additional *C. gloeosporioides* species complex isolates, which is shown in Fig. 5C. A 100 bp or 1 kb ladder was used to provide size estimates. See Supplementary Table S5 for details of the strains used.

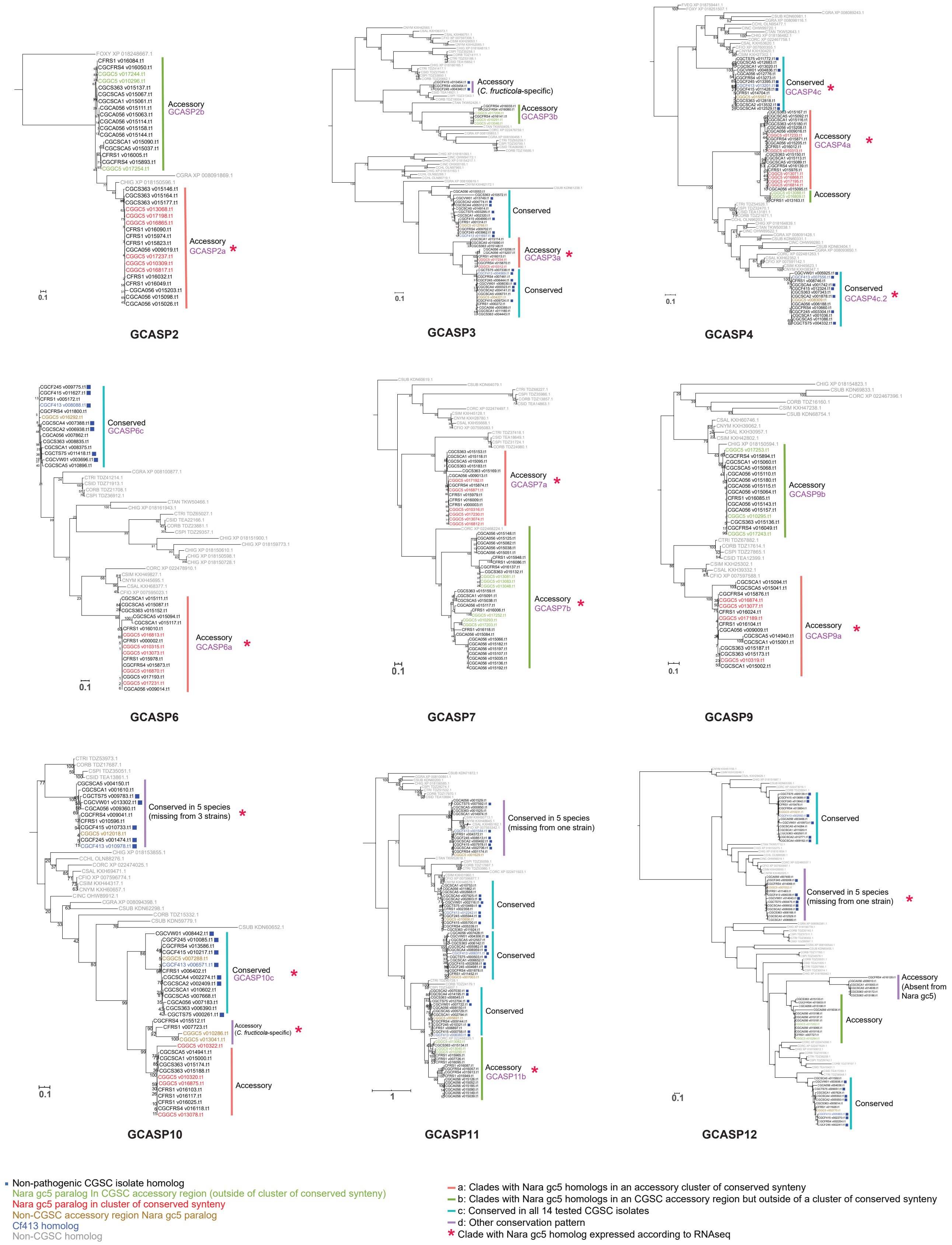


**Supplementary Fig. S11.** Maximum likelihood phylogenies of GCASP homologs with paralogs outside of accessory clusters of conserved synteny in *Colletotrichum fructicola* Nara gc5. GCASP1, 5 and 8 are not included as these sequences are only present in accessory clusters of conserved synteny. Values at nodes are percentages of support of 1000 bootstrap replicates. Purple labels: sequences used to design primers used for qPCR analysis (Supplementary Table S7 and Supplementary Fig. S12). The branch length of CSAL KXH65162.1 in GCASP11 is shortened 10-fold.

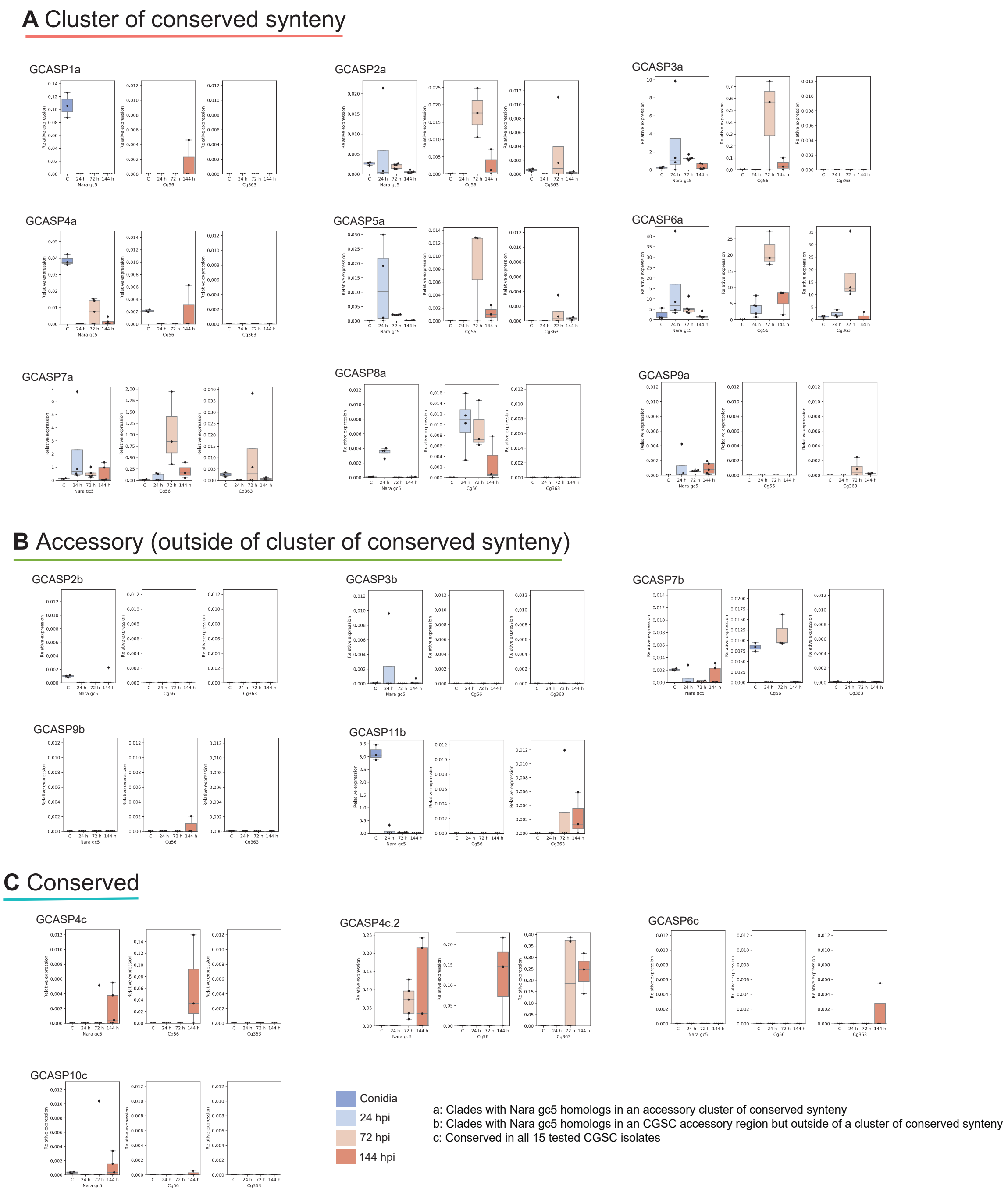


**Supplementary Fig. S12.** Quantitative PCR of selected GCASP homologs in *Colletotrichum fructicola* (*Cf*) Nara gc5, *C. aenigma* Cg56 and *C. siamense* Cg363 designed based on *Cf* Nara gc5 sequences in (A) accessory clusters of conserved synteny, (B) CGSC accessory regions and (C) regions that are conserved in all 14 sequenced CGSC strains. Expression levels relative to elongation factor 1 (*CfEF*) were assessed in conidia and in *F. vesca* infected leaves at 24 , 72 and 144 hours post-inoculation (hpi). Data from 3-5 replicates per time point are shown with a boxplot showing the distributions of gene expression. Only sequences with at least one time point showing expression in any of the three strains are shown here.

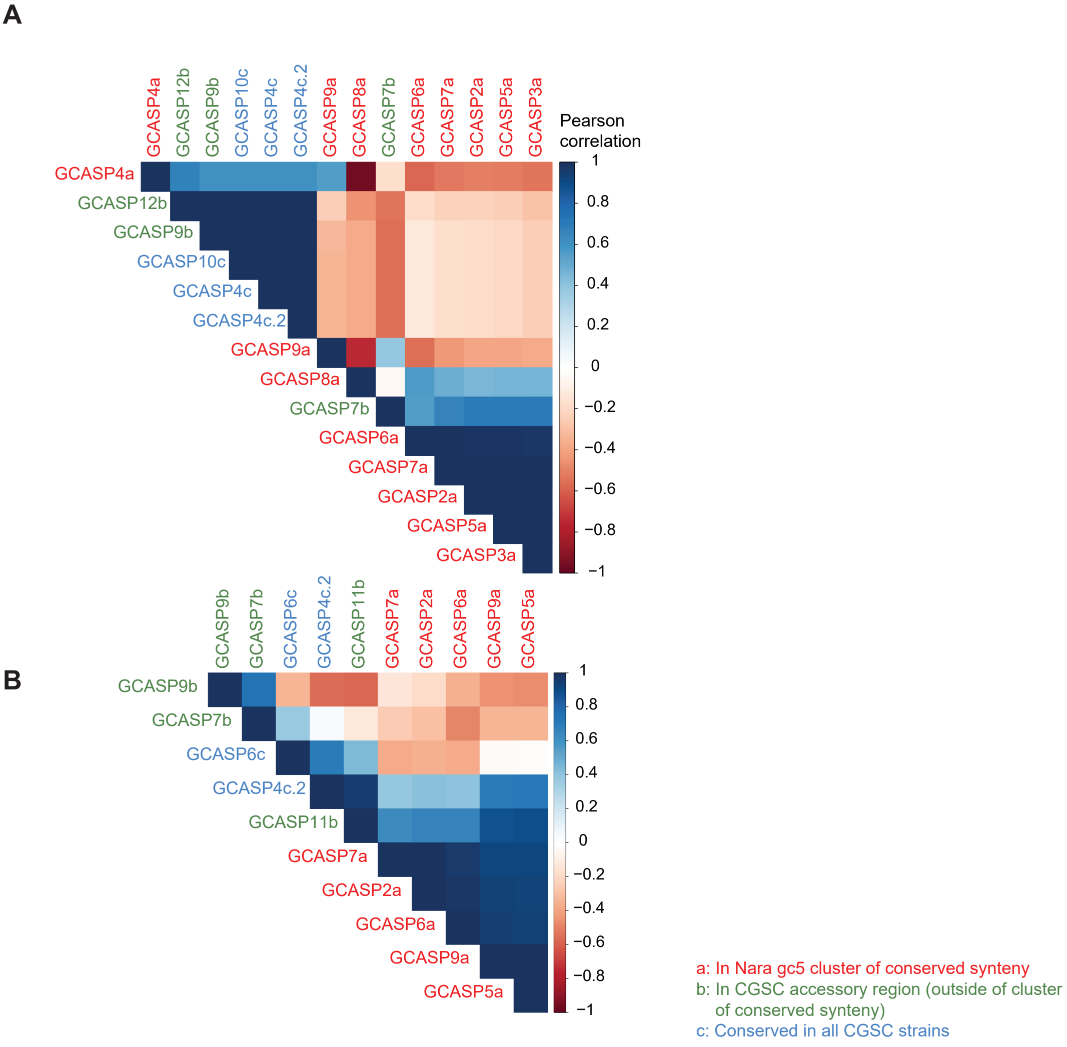


**Supplementary Fig. S13.** Correlation plots to visualize the correlation of GCASP homolog genes expression in (A) *Colletotrichum aenigma* Cg56 and (B) *C. siamense* Cg363. Mean relative expression levels in each strain were scaled within each primer set and then pairwise correlation scores were calculated.
